# YY1 relieves p300 autoinhibition to promote histone acetylation in advanced prostate cancer, enhancing the oncogenic signaling of Androgen Receptor Splice Variant 7

**DOI:** 10.1101/2025.08.04.668477

**Authors:** Chenxi Xu, Damu Wu, Bo Pan, Xiaoxiao Ren, Samuel G. Mackintosh, Kanishk Jain, Natarajan V Bhanu, Ricky D. Edmondson, Xin Liu, Benjamin A Garcia, Alan J. Tackett, Jian Jin, Gang Greg Wang, Ling Cai

**Author notes:** Correspondence: G.G.W. and L.C.

## Abstract

Genetic and epigenetic aberrations often act in concert to establish oncogenic transcriptomic programs in aggressive cancers. For example, the development of castration-resistant prostate cancer (CRPC), an advanced prostate cancer form, is closely associated with over-expression and/or hyper-activation of transcription factors (TFs) such as Androgen Receptor (AR) and Yin Yang 1 (YY1), as well as p300, a prominent histone acetyltransferase. How exactly these cancer-related lesions are coordinated to generate a malignant cell state remains elusive. Here, we demonstrate that YY1, which is frequently over-expressed in advanced prostate cancers, allosterically stimulates the acetyltransferase activity of p300 *in cis*, leading to the globally elevated acetylation of histone H3 lysine 18 and 27 (H3K18ac and H3K27ac). Mechanistically, YY1’s N-terminal activation domain (AD) directly interacts with p300’s TAZ2 domain, relieving the autoinhibition of p300 to facilitate substrate acetylation. Our integrated genome-wide mapping and transcriptomic studies reveal significant co-localization of genomic binding sites of YY1, androgen receptor splice variant 7 (AR-V7, a constitutively active form of AR) and p300 in CRPC cells, where the YY1-mediated p300 activation and resultant histone acetylation increases promote the oncogenic gene-expression programs downstream of YY1 and AR/AR-V7. Both *in vitro* and *in vivo* functional assays demonstrate a critical requirement of the above signaling for the advanced disease progression and drug resistance seen in CRPC. Altogether, this study uncovers that YY1 acts to alleviate p300’s autoinhibition at target genes co-bound by oncogenic TFs (YY1 and/or AR/AR-V7) in CRPC, thereby sustaining tumorigenicity. Additionally, the blockade of YY1-mediated gene activation resensitizes CRPC to treatment to the clinic anti-AR agent (enzalutamide), which provides a rationale for overcoming the therapeutic resistance often seen in advanced prostate cancers.

## Introduction

Prostate cancer is one of the leading causes of cancer-related deaths among the men world-wide^1^. Androgen receptor (AR), a lineage-specific transcription factor (TF), plays a central role in the development and advanced progression of prostate cancer^2–4^. Despite the development and clinical usage of the highly specific and highly potent anti-androgen agents, prostate cancers remain incurable due to therapeutic resistance. Indeed, most of the prostate cancer patients receiving the anti-androgen treatments eventually develop the so-called castration-resistant prostate cancer (CRPC), a terminal form of this disease. Previous studies have linked CRPC pathogenesis to AR dysregulations, which include the AR gene amplification and activating mutations^2,5,6^, the expression of a constitutively active AR splice variant such as androgen receptor splice variant 7 (AR-V7, which lacks AR’s ligand-binding domain)^7–10^, tumor-derived androgen production^11^, over-expression of AR’s coactivators, amongst others. Aside of AR dysregulations, Yin Yang 1 (YY1), another TF frequently exhibiting an over-expressed pattern in prostate cancer^12^, emerges as a critical player in CRPC as well. Furthermore, more aggressive features seen with advanced prostate cancers were also found to be correlated with hyper-activation of p300 (also known as EP300 and KAT3B), a prominent histone acetyltransferase and transcriptional coactivator^13–15^, indicative of the involvement of epigenetic lesions during the disease progression. Indeed, the elevated levels and/or functionalities of AR^7–10^, YY1^12^ and p300^15^ were all linked to the advanced progression of prostate cancers. How exactly these genetic and epigenetic lesions act in concert to establish the oncogenic transcriptomic programs in CRPC, a currently incurable disease, remain poorly understood. A better understanding of the above events shall help to develop the more effective treatment of the affected patients.

In cells, the enzymatic activity of p300 is subject to exquisite regulation, such as autoinhibition, the TF-induced trans-autoacetylation and activation, as well as protein condensation^16–20^. P300 contains multiple conserved functional modules, which include cysteine–histidine-rich region 1 to 3 (CH1 to CH3), kinase inducible domain-interacting (KIX) domain, bromodomain (BD) and histone acetyltransferase (HAT) domain (**Supplementary Fig. S1A**). P300’s CH3 domain comprises two zinc-binding motifs, namely, ZZ-type zinc finger (ZZ) and transcriptional adapter zinc-binding domain 2 (TAZ2). While the p300 ZZ domain serves as a reader of histone H3’s N-terminus, allowing selective acetylation of H3K18 and H3K27 (H3K18ac and H3K27ac)^21,22^, TAZ2 is increasingly appreciated as a critical protein-protein interaction interface for mediating direct association with TFs (such as p53 and IRF3) and chromatin factors (such as BRD4-NUT)^17,18,20,23^. Binding of TAZ2 to the trans-activation domain (AD) of TFs was reported to cause conformational change of the p300 HAT domain from a closed state to open, enhancing histone acetylation^17,18,20,23^. The CH1 of p300 was reported to bind TFs as well^24–26^. While the loss-of-function somatic mutation of p300 is recurrent in B cell malignancies^27,28^, p300 is frequently overexpressed in solid cancers, notably advanced prostate cancers, and correlates with poor clinical outcomes^15^, speaking to complex context-dependent roles for p300. How exactly the p300 activity and related histone acetylation landscape is regulated in prostate cancer needs to be further elucidated.

In this study, we report that YY1, but not other tested prostate cancer-related TFs (such as AR and FOXA1), is required for elevating the p300-mediated H3K27ac and H3K18ac at the global level in CRPC. Mechanistically, YY1’s N-terminal AD directly binds p300’s TAZ2 domain to activate p300’s HAT activity. Also, we took an integrative genomic analysis approach to investigate the role of the above YY1:p300-mediated signaling in the regulation of the histone acetylation landscape, oncogenic gene-expression program, and CRPC tumorigenesis and drug resistance. YY1-overexpressed CRPC cells are more sensitive to p300 inhibition, offering an attractive anti-tumor strategy. Collectively, our observations highlight a crucial involvement of the crosstalk between cancer-associated TFs and chromatin factors for generating the aggressive cancer phenotypes, suggesting a therapeutic strategy for treating the otherwise drug-resistant patients.

## Results

### YY1, a frequently overexpressed TF in prostate cancer, elevates the p300-mediated histone acetylation globally

How exactly the enzymatic activity of p300 is regulated in advanced PCa is not fully characterized. To identify factors potentially involved in the p300 activity regulation, we first conducted an unbiased proximity-dependent biotin identification (BioID) approach to investigate the p300 interactome in 22Rv1 cells, a commonly used cell model of CRPC (**Supplementary Fig. S1B**). Following the p300 BioID, mass spectrometry identified the known p300-interactors such as HCFC1^29^, BRD4^30^ and CHD4^31^ (**Fig. 1A**). Additionally, YY1 was discovered as one of the top enriched hits (**Fig. 1A**). Previously, we have shown that YY1 plays a crucial role during the tumorigenesis of advanced prostate cancer^32^. Co-Immunoprecipitation (Co-IP) validated the interaction between p300 and YY1, together with BRD4 (**Fig. 1B**). We also observed the interaction between p300 and AR, both the full-length form of AR (AR-FL) and AR-V7 (**Fig. 1B**), consistent with a previous report^33^. Elevated levels and/or functionalities of YY1 and p300 were suggested to underlie the advanced progression of prostate cancers^12,15^. The identified YY1:p300 interaction indicates a role for YY1 in modulating the p300 activity and histone acetylation landscape in CRPC cells. Towards this end, we conducted YY1 depletion in both 22Rv1 and VCaP CRPC cells (**Fig. 1C, top: YY1**), followed by quantitative mass spectrometric profiling of total histones (**Fig. 1D**). This analysis revealed the consistent decrease of H3K27 acetylation (H3K27ac) in both cell types upon YY1 depletion (**Fig. 1D**). By western blot (WB) of a panel of individual histone marks, we verified that YY1 depletion led to the significantly reduced H3K27ac and H3K18ac in 22Rv1 and VCaP cells, whereas global acetylation levels at the other tested histone lysine sites, such as H3K9, H3K14, H3K36, H3K56 and H4, all remained largely unaffected (**Fig. 1C**). H3K27ac and H3K18ac are the major histone acetylations deposited by p300^34^. To further validate the impact of YY1 on histone acetylation and rule out any potential siRNA-related off-target effect, we additionally employed an auxin-inducible degron version 2 (AID2) system^35^ for achieving rapid and efficient degradation of YY1 (**Supplementary Fig. S1C**). Here, we first knocked in a mini-AID (mAID) and 3×Flag tag in-frame to the C-terminus of endogenous YY1 (**Supplementary Fig. S1D**) and verified validity of such YY1-mAID alleles in the engineered 22Rv1 cells using both sanger sequencing and WB (**Supplementary Fig. S1D-E**). YY1-mAID fusion was rapidly degraded upon the addition of 5-Ph-IAA, the ligand of mAID, in a time-dependent manner (**Supplementary Fig. S1F**), which caused the expected suppression of 22Rv1 cell growth (**Supplementary Fig. S1G, blue vs. red and green**). As a control, fusion of mAID to YY1 *per se* did not affect cell proliferation (**Supplementary Fig. S1G, green vs. gray**). Following YY1 degradation, we observed a marked reduction in global H3K27ac and the expression of PFKP, a known target of YY1^32^ (**Fig. 1E**). YY1 loss did not alter the level of p300 protein (**Fig. 1C**, see panels of p300); thus, the global decrease of H3K27ac and H3K18ac is likely due to a change in p300’s enzymatic activity upon the depletion of YY1, which hasn’t been reported before.

**Figure 1.**
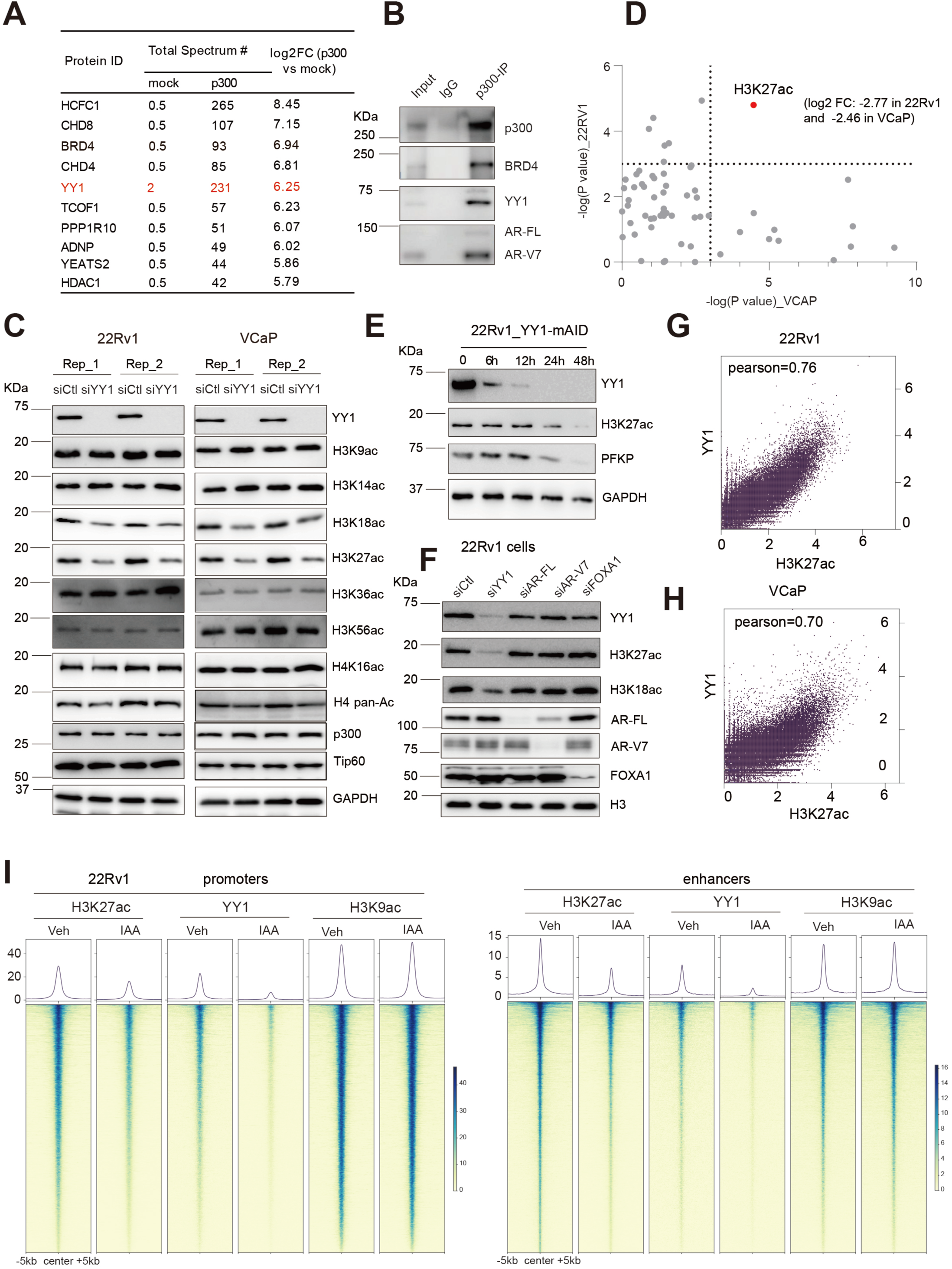
YY1, and not other tested oncogenic TFs (AR, AR-V7 or FOXA1), enhances the p300-mediated histone acetylation globally in CRPC. (A) BioID identified the p300-associated proteins in 22Rv1 cells, with hits ranked by the fold change (FC) of normalized spectral abundance factor (NSAF) relative to empty vector (EV) control. (B) Co-immunoprecipitation (Co-IP) in 22Rv1 cells for interaction between endogenous p300 protein and the identified potential partners, YY1, BRD4 and AR (either full-length AR [AR-FL] or AR-V7). IgG was used as a control. (C) WB of the indicated protein in 22Rv1 (left) and VCaP (right) cells 48 hours post-transfection of non-specific control siRNA (siCtrl) or YY1-targeting siRNA (in duplicate). (D) Unbiased mass spectrometry-based histone modification quantification using total histones isolated from 22Rv1 (y-axis) and VCAP cells (x-axis) with YY1 depleted, compared to mock treated (n= 3 biological replicates per group). Plotted are -log converted values of *P* value for change of each histone modification (YY1-depleted versus mock), with FC values for H3K27ac (red) labeled aside. (E) WB of the indicated protein in 22Rv1 cells carrying the engineered YY1-mAID knock-in (KI) alleles, treated with 2.5 µM of 5-Ph-IAA for 0, 6, 12, 24 or 48 hours. GAPDH was used as a loading control. (F) WB of the indicated protein in 22Rv1 cells, transfected with control siRNA (siCtrl, lane 1) or those targeting either YY1, AR-FL, AR-V7 or FOXA1 (lanes 2-5). **(G-H)** Pearson correlation plot using the YY1 and H3K27ac CUT&Tag profiles of 22Rv1 (**G**) and VCaP (**H**) cells. **(I)** Heatmap showing the H3K27ac, YY1 and H3K9ac CUT&Tag read densities at the promoters (left) and enhancers (right) in 22Rv1 cells, treated with either vehicle (Veh) or 2.5 µM of 5-Ph-IAA (IAA) for 48h in. The YY1 peaks were sorted based on read densities, and then the signals of all factors were shown ±5 kb from the centers of YY1 peaks.

Interestingly, the histone acetylation enhancement effect appeared specific to YY1, because depletion of other prominent oncogenic TFs in prostate cancer, such as full-length AR (AR-FL), the constitutively active AR-V7 or FOXA1 (an AR-associated pioneer factor^36^), did not affect the H3K18ac or H3K27ac levels (**Fig. 1F**). Using the available p300 enzymatic inhibitor, A485^17^, we verified the requirement of p300 for specific acetylation of H3K27 and H3K18, but not H3K9 or H3K14, in CRPC cells (**Supplementary Fig. S1H).** Additionally, treatment of same cells with enzalutamide, a potent anti-AR agent used in the clinic, did not alter the H3K27ac or H3K18ac levels (**Supplementary Fig. S1H)**, consistent with the results upon AR depletion (**Fig. 1F**).

Next, we aimed to map out the genomic regions where H3K27ac deposition might be affected by YY1 loss. Here, we conducted Cleavage Under Targets and Tagmentation (CUT&Tag) of YY1 and H3K27ac in two independent CRPC models (22Rv1 and VCaP cells), either mock-treated or YY1 depleted. The results revealed a striking co-localization between YY1 binding and H3K27ac genome-wide in both CRPC cells (**Fig. 1G-H**). Upon YY1 depletion, we consistently observed the markedly decreased H3K27ac at the YY1-bound genomic sites, regardless of promoters or enhancers, following either the mAID-mediated rapid degradation in 22Rv1_YY1-mAID cells (**Fig. 1I**) or shRNA-mediated YY1 knockdown (KD) in VCaP cells (**Supplementary Fig. S1I**). Meanwhile, we conducted CUT&Tag for H3K9ac and found that YY1 loss did not affect the genome-wide H3K9ac occupancies, which contrasts the dramatic change seen with H3K27ac (**Fig. 1I**, see panels of H3K9ac versus H3K27ac) and agrees with WB data (**Fig. 1C**, see panels of H3K9ac versus H3K27ac). The requirement of YY1 for deposition of H3K27ac, but not H3K9ac, was exemplified at key prostate cancer biomarker genes such as KLK3 (aka, prostate-specific antigen or PSA) and KLK2 (**Supplementary Fig. S1J**).

Altogether, there exists an exquisite requirement of YY1 for global deposition of the classic histone acetylations associated with p300’s HAT activity (namely, H3K27ac and H3K18ac) in CRPC cells.

### YY1 increases p300’s HAT activity through its N-terminal domain

Having demonstrated a striking enhancement effect by YY1 on the p300-mediated histone acetylations, we further aimed to investigate their functional and physical association in CRPC. Over-expression of YY1 and p300 has been reported in advanced prostate cancers^12,15^. Interestingly, there exists a positive correlation between the YY1 and p300 expression levels in independent clinical samples of patients with metastatic CRPCs, as shown in in the Abida et al., PNAS 2019^4^ and Robinson et al., Cell 2015^37^ datasets via cBioPortal (**Fig. 2A and Supplementary Fig. S2A**), suggesting that they may cooperate in promoting the progression of this lethal disease. Using a panel of prostate cancer patient samples and paired benign tissues, we confirmed that YY1 and p300 are often upregulated in prostate cancer relative to respective benign tissue controls, and additionally, observed that prostate cancer samples with over-expression of both YY1 and p300 tend to have a globally elevated level of H3K27ac (**Fig. 2B**). Consistent with YY1 being enriched in the p300 interactome (**Fig. 1A**), immunofluorescence (IF) readily detected the colocalization of YY1, p300 and H3K27ac in 22Rv1 and VCaP cells (**Fig. 2C and Supplementary Fig. S2B**), which agrees with the positive correlation, at a genome-wide scale, seen between YY1 and p300 binding sites (**Fig. 2D**), or those of YY1 and H3K27ac (**Fig. 1**). To determine whether or not YY1 loss affected the p300 chromatin association, we performed the cell fractionation assay—here, we found that YY1 loss caused the H3K27ac decrease while having no effect on the level of chromatin-bound p300 (**Fig. 2E**, lane 4 versus 3). To further assess if YY1 influences p300’s chromatin targeting in a genome-wide scale, we performed CUT&Tag of p300 in 22Rv1 cells before and after inducing YY1 degradation—again, YY1 depletion did not cause significant change in global binding patterns of p300 (**Fig. 2F**), suggesting that YY1 might instead regulate the HAT activity of p300 at their co-bound genomic sites.

**Figure 2.**
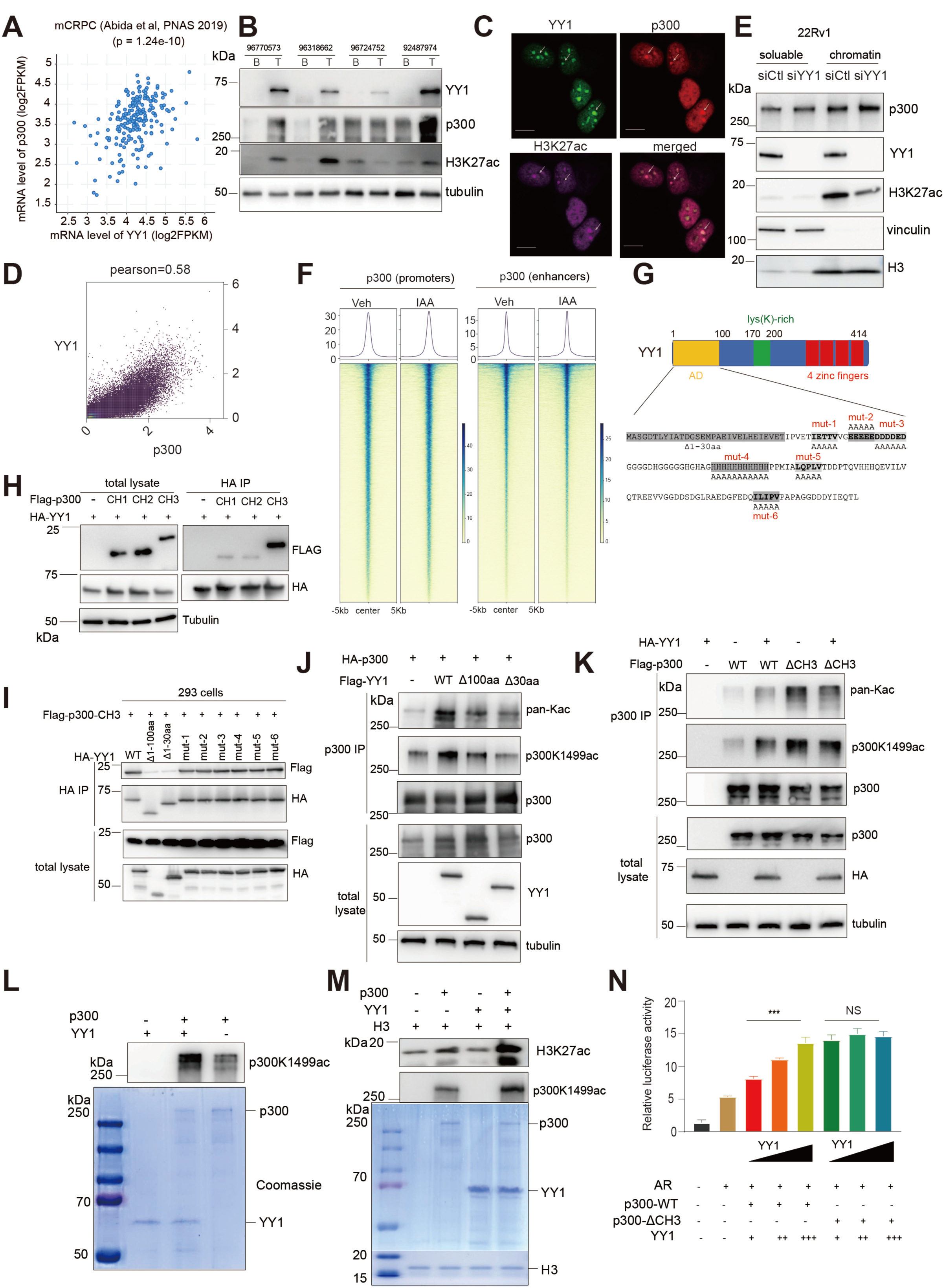
YY1’s N-terminus interacts with p300’s CH3 module, relieving p300 autoinhibition to promote p300 trans-autoacetylation and substrate acetylation for gene activation. (A) Correlation plots using the mRNA expression levels of YY1 (x-axis) and p300 (y-axis) in the metastatic castration-resistant prostate cancer (mCRPC) samples from the Abida et al (2019) dataset (generated via cBioPortal; n = 444, Pearson correlation, p = 1.24e– 10). (B) WB for H3K27ac, p300 and YY1 using total protein lysates of the paired benign (B) and tumor (T) tissues from five CPRC patients. Tubulin acts as a loading control. (C) Immunofluorescence (IF) of the indicated protein in 22Rv1 cells. Representative results from three independent experiments. Scale bar, 10 μm. **(D)** Pearson correlation plot using the YY1 and p300 CUT&Tag profiles in 22Rv1 cells. **(E)** WB of p300, either nucleoplasmic (left) or chromatin-bound (right), in 22Rv1 cells after depleting YY1 (siYY1) versus mock control (siCtl). Vinculin and histone H3 served as cell fractionation control. **(F)** Heatmap of the p300 CUT&Tag read densities at the promoter (left) and enhancer (right) peaks in 22Rv1 cells, treated with either vehicle (Veh) or 2.5 µM of 5-Ph-IAA (IAA) for 48h in. **(G)** Top: Schematic diagram of the YY1 protein highlighting its functional domains. The trans-activation domain (AD), a lysine-rich region and zinc fingers are shown in yellow, green and red, respectively. Bottom: a set of N-terminal truncation and alanine substitution scanning mutants used to assess the region for mediating the YY1–p300 interaction. **(H-I)** Co-IP for interaction in 293 cells using the exogenously expressed HA-tagged YY1, either WT **(H)** or a set of the indicated truncation and substitution mutants of YY1 (shown in **G: bottom panel**), and individual p300^CH^ module, either one of three CHs (**H**; CH1, CH2 or CH3) or CH3 alone (**I**). **(J-K)** Assessing p300 trans-autoacetylation levels (WB with both anti-p300K1499ac and pan-Kac antibodies) post-IP of p300 (**J-K: top panel**) from 293 cells co-transfected with tagged p300, either WT **(J)** or the indicated CH3 internal deletion form **(K;** p300_ΔCH3 versus WT), and YY1, either WT or truncated (YY1_Δ1-100aa or YY1_Δ1-30aa versus WT). Whole-cell lysates were collected and used as a control (**J-K: bottom panel**). **(L-M)** *In vitro* acetylation, in the presence of acetyl-CoA, to assess the p300 trans-autoacetylation **(L-M)** and substrate (H3) acetylation levels **(M)** using the recombinant full-length p300 and YY1 protein, without **(L)** or with addition **(M)** of the recombinant H3 protein. Reaction products were analyzed by SDS–PAGE followed by WB with an antibody against p300K1499ac or H3K27ac (top panels), with Coomassie blue staining of input shown in bottom panels. **(N)** Summary of reporter activation assays in HEK293 cells, co-transfected with an androgen response element-luciferase reporter (ARE-Luc) and AR, along with p300 (either full-length WT p300 or p300_ΔCH3) and an increasing amount of YY1. Relative luminescence was normalized to the renilla internal controls and to mock-transfected cells (n=3 independent experiments; shown as mean ± SD). \*\*\**P* < 0.001.

While the C-terminal region of YY1 has four conserved C2H2 zinc fingers that bind DNA, its N-terminal segment is largely intrinsically disordered and contains a trans-activation domain (AD) enriched with motifs of acidic residues, histidine and hydrophobic residues^38^ (**Fig. 2G-top**), some of which was reported to promote condensate formation^39^. TFs, often via their ADs, have been reported to interact with p300 through its CH1, CH2 and CH3 modules^24–26^. To evaluate whether YY1 also interacts with p300 via these CH regions, we first performed co-IP using full-length YY1 and individual p300 CH1-3 segment and found that YY1 strongly bound to p300’s CH3 (p300^CH3^) and barely associated with CH1 and CH2 domains (**Fig. 2H**). To test if the N-terminal segment of YY1 mediates p300 interaction, we employed p300^CH3^ for co-IP with YY1, either WT or various truncated and point mutants of its N-terminal region. Compared to wildtype (WT) YY1 control, deletion of the first 1-100 amino acids (YY1_Δ1-100aa) or the first 1-30 amino acids (YY1_Δ1-30aa) significantly reduced the p300^CH3^ association (**Fig. 2I**, see lanes 1-3; **Supplementary Fig. S2C**); meanwhile, none of the tested alanine substitution mutants (**Fig. 2I**) that were designed to disrupt the acidic residue cluster (mut-2 and mut-3), the histidine cluster (mut-4) or hydrophobic residue motifs after the first 30 amino acids (mut-1, mut-5 and mut-6) affected YY1 binding to p300^CH3^ (**Fig. 2I**). These data highlight a critical role of YY1’s first 30 amino acids (YY1^1-30aa^) in mediating the p300 binding.

Acetylation of p300 at its lysine 1499 residue (p300K1499ac) was reported as an indicator of p300 enzymatic activation following the TF:p300 interaction and p300 trans-autoacetylation^17^. To examine putative effect of YY1 on p300’s enzymatic activity, we ectopically expressed YY1 with HA-tagged full-length p300 into 293 cells. Such p300 protein was pulled down from cells using tag-specific antibody and then probed with the antibody recognizing p300K1499ac or acetyl-lysine (pan-Kac). Compared to mock, we readily detected that WT YY1, but not its truncated version that failed to bind p300 (Δ1-100aa or Δ1-30aa, as tested in **Fig. 2I**), increased p300 activity in cells (assessed by autoacetylation at p300K1499ac; **Fig. 2J**, lane 2 versus 3-4). Likewise, we co-transduced p300, either WT or a CH3-deleted mutant (p300-ΔCH3), together with full-length YY1 into 293 cells, which revealed that YY1 strongly enhanced acetylation of WT p300, but not its ΔCH3 form (**Fig. 2K**, lane 3 versus 2 and lane 5 versus 4). As expected, there existed a clearly enhanced autoacetylation of p300-ΔCH3, relative to WT p300, in the absence of YY1 (**Fig. 2K**, lane 4 versus 2), consistent with the role for CH3 in mediating p300 autoinhibition^17,18^. As a negative control, no acetylated-p300 was detected using the immunoprecipitated p300 form lacking HAT domain (p300-ΔHAT), when compared to WT p300 controls (**Supplementary Fig. S2D**, lanes 4-5 versus 2-3). These results further support that the YY1 can directly increase the HAT activity of p300 *per se*. To further substantiate the YY1-mediated p300 activity stimulation, we employed the *in vitro* acetylation assays with the recombinant protein of full-length p300 and YY1. In the presence of acetyl-CoA (a cofactor of p300 and the donor of acetyl group), YY1 significantly enhanced p300 autoacetylation (p300K1499ac) *in vitro* (**Fig. 2L**, lane 2 versus 3). In a separate *in vitro* HAT assay, we additionally included a H3K27-containing histone peptide and detected H3K27ac by recombinant p300 in the presence of YY1 (**Fig. 2M**, lane 4 versus 2).

To further demonstrate that YY1 can indeed stimulate p300’s HAT activity through its interaction with p300^CH3^ domain in cells, we constituted the complex with a classic luciferase reporter carrying an upstream androgen responsive element (ARE-Luc). In the presence of WT p300 and AR, YY1 dose-dependently elevated transcription from the reporter (**Fig. 2N**, columns 3-5 versus 1-2); however, in the context of p300-ΔCH3, the reporter activity was generally higher than what was seen in the presence of WT p300, and additionally, it did not respond to the level of YY1 (**Fig. 2N**, see last three columns).

Taken together, the N-terminal AD of YY1 binds the CH3 domain of p300 to relieves its autoinhibition, thereby stimulating p300’s HAT activity *in vitro* and in cells. Because YY1 and p300 were reported to be overexpressed in advanced prostate cancers and correlated with poorer outcomes^15,32^, such a regulatory axis involving YY1^AD^:p300^CH3^ is clinically relevant.

### Structural modeling of the p300^TAZ2^:YY1^AD^ interaction

Having demonstrated a physical and functional interaction between YY1’s N-terminal segment (YY1^1-30aa^) and p300^CH3^, we then aimed to gain the molecular details underlying this interaction. Here, we employed AlphaFold2 to conduct structural modeling studies. The two most confident models revealed that YY1^1-30aa^ interacts with the TAZ2, but not ZZ, module of p300^CH3^: in both models, p300’s TAZ2 domain (p300^TAZ2^) adopts a consistent and rigid conformation whereas YY1^1-30aa^ displays notable conformational flexibility, primarily manifested as distinct orientations relative to p300^TAZ2^ (**Supplementary** Fig. 3A-B). In both predicted complexes, YY1^1-30aa^ and p300^TAZ2^ form extensive interactions through two sets of interaction motifs, namely, acidic residues (E19, E22 and E25; **Fig. 3A-B)** and hydrophobic residues (I20 and L23; **Fig. 3C-D**). Specifically, electrostatic surface analysis revealed that three acidic residues of YY1 (E19, E22 and E25) align with a positively charged surface of p300^TAZ2^ (**Fig. 3A-B**), while I20 and L23 of YY1 interact with a hydrophobic pocket on p300^TAZ2^ (**Fig. 3C-D**). To confirm this predicted interaction, we conducted GST pull-down and found that YY1^1-30aa^ directly binds p300^TAZ2^ (**Fig. 3E**), which was further supported by isothermal titration calorimetry (ITC) assay (**Supplementary Fig. S3C**). Notably, this YY1 N-terminal sequence is highly conserved across metazoan species (**Fig. 3F**). To investigate significance of the above putative protein-protein interaction interfaces, we substituted YY1’s E19, E22 and E25 with arginine in one mutant while mutating I20 and L23 to serine in the other mutant (hereafter referred to respectively as YY1^3E-3R^ and YY1^IL-SS^; **Fig. 3F**). When compared to WT, both mutants exhibited a significantly reduced binding to p300^TAZ2^ (**Fig. 3G**, lane 1 versus 3-4). When we combined both sets of mutations together (hereafter referred to as YY1^5aa-mut^; **Fig. 3F**), the binding between full-length YY1 and p300^TAZ2^ was almost completely abolished, resembling what was observed for the Δ1-30aa truncated version of YY1 (**Fig. 3G**, lane 5 versus 1-2). Furthermore, the capability of YY1 in enhancing p300 autoacetylation was significantly impaired by the YY1^5aa-mut^ compound mutant (**Fig. 3H**). Altogether, the N-terminal region of YY1 (YY1^1-30aa^) directly interacts with p300^TAZ2^, leading to p300 trans-autoacetylation and activation.

**Figure 3.**
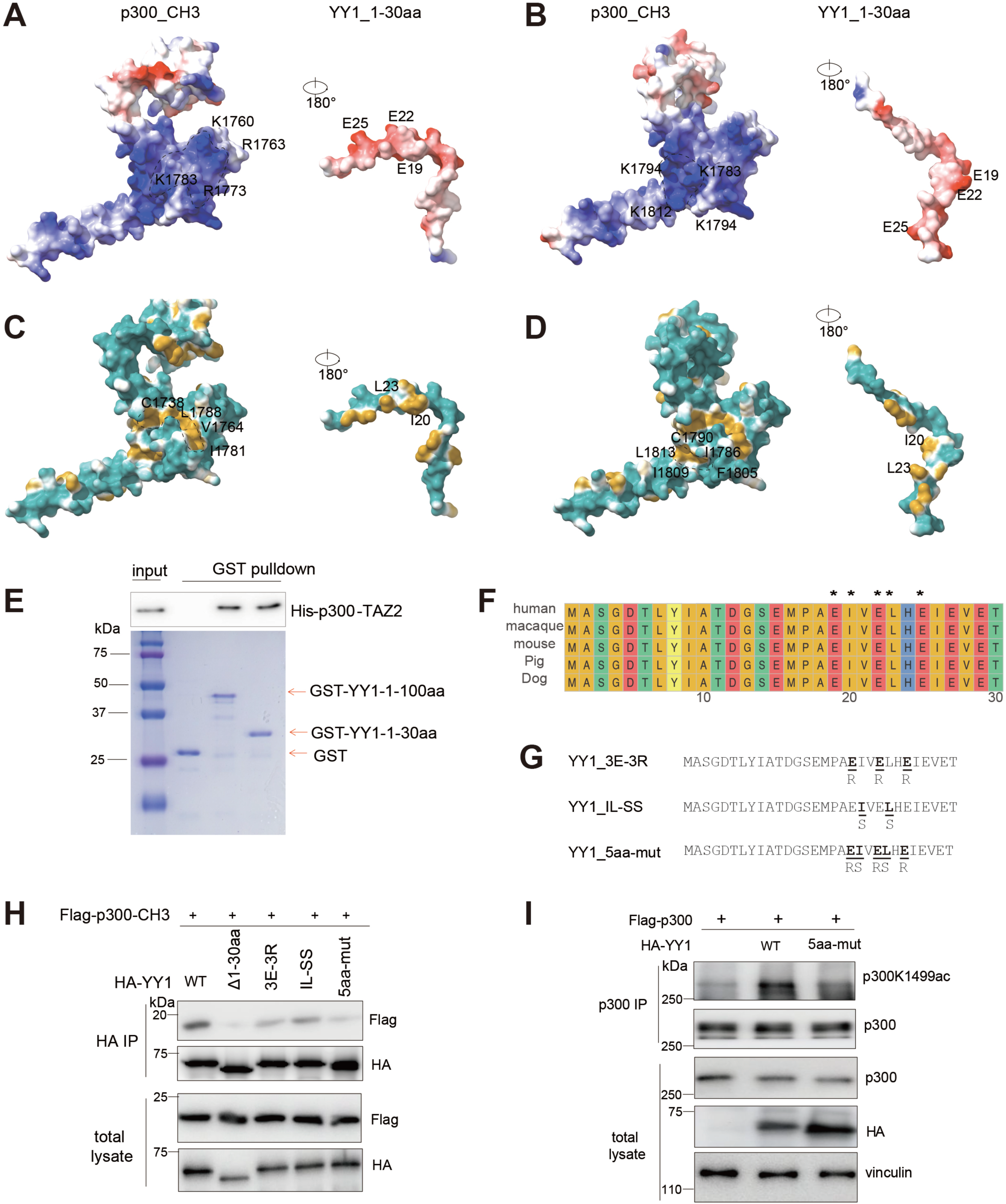
Structural and biochemical analyses of the p300^CH3^:YY1^AD^ interaction. (A–D) Predicted electrostatic interaction **(A–B)** and hydrophobic interaction **(C–D)** interfaces between YY1’s N-terminus (1–30aa) and p300^CH3^ in the two top-ranked models based on AlphaFold2 structural, either Model 1 (**A, C**) or Model 2 (**B, D**), illustrating potential electrostatic and hydrophobic contacts as well as conformational differences between the two models. **(E)** GST pulldown using the GST fusion protein, which contained either GST alone or GST fused to YY1_1-100aa or YY1_1-30aa, and the cell lysate containing His6-tagged p300-TAZ2, followed by anti-His-tag WB (top). **(F)** Sequence alignment of the N-terminal 1–30 amino acids of YY1 from human, macaque, mouse, pig, and dog. **(G)** Schematic of the three used YY1^AD^ mutants, including a three-acidic-residues mutant (E19R/E22R/E25R; termed as 3E-3R), a two-hydrophobic-residues mutant (I20S/L23S; termed as IL-SS), and a compound mutant harboring 3E-3R and IL-SS both (termed as 5aa-mut). **(H-I)** Co-IP for interaction between the exogenously expressed Flag-tagged p300, either p300^CH3^ alone **(H)** or full-length WT p300 **(I)**, and the indicated HA-tagged YY1 (WT versus mutant such as 5aa-mut or others) in HEK293 cells. Whole-cell lysates were used for IP with anti-HA **(H)** or anti-p300 antibody **(I)**. Equal amounts of IP (top) and whole-cell lysate samples (bottom; as a loading control) were subject for WB of p300 or YY1, with IP samples in **I** additionally probed with the p300K1499ac antibody.

### YY1^1-30aa^ is essential for the YY1-induced oncogene activation in CRPC

To further assess the gene-regulatory role of YY1^1-30aa^ in CRPC cells, we performed RNA-seq using the YY1-depleted CRPC cells reconstituted with YY1, either WT or the YY1^1-30aa^-truncated mutant (YY1^Δ1–30aa^). Differentially expressed genes (DEGs) identified after the rescued expression of YY1^Δ1–30aa^ relative to WT YY1 in 22Rv1 cells (**Fig. 4A**) and VCaP cells (**Supplementary Fig. S4A**) showed more transcripts to be downregulated than upregulated. Over half of the downregulated DEGs due to YY1^1-30aa^ truncation overlapped those downregulated by YY1 KD in 22Rv1 cells (**Fig. 4B-C, Supplementary Fig. S4B**). These results support a gene-activating role for YY1^1-30aa^. Gene Ontology (GO) and Gene Set Enrichment Analysis (GSEA) further revealed that the transcripts showing the YY1^1-30aa^ dependence for upregulation in 22Rv1 cells were primarily associated with hormone signaling, AR and AR-V7-related gene pathways and the YY1 targets (**Fig. 4D-E** and **Supplementary Fig. S4C**), as well as the proliferative transcripts such as cell cycle genes, E2F targets, MYC targets, and prostate cancer oncogenes in 22Rv1 cells (**Fig. 4F**). Similar results were observed in another CRPC model, VCaP cells (**Supplementary Fig. S4D-F**). Moreover, analyses using the public transcriptomic datasets of patient samples^40–42^ showed a strong correlation between the expression of YY1 and p300 with the AR/AR-V7 signature genes (**Fig. 4G-H**). RT-qPCR not only confirmed that YY1 promotes the activation of the select transcripts known to be the AR/AR-V7 targets (namely, KLK2, KLK3, PMEPA1, SREBF1, ECE1 and SCN8A; **Fig. 4I**, see vehicle [Veh] vs. IAA), but also demonstrated that YY1^1-30aa^ (especially E19, E22, E25, I20 and L23) is essential for activating these transcripts in 22Rv1 cells (**Fig. 4I**, see YY1_WT vs. YY1^Δ1–30aa^ and YY1^5aa-mut^). Collectively, YY1^1-30aa^ plays crucial roles in enhancing oncogenic programs of CRPC, including the AR/AR-V7 signaling.

**Figure 4.**
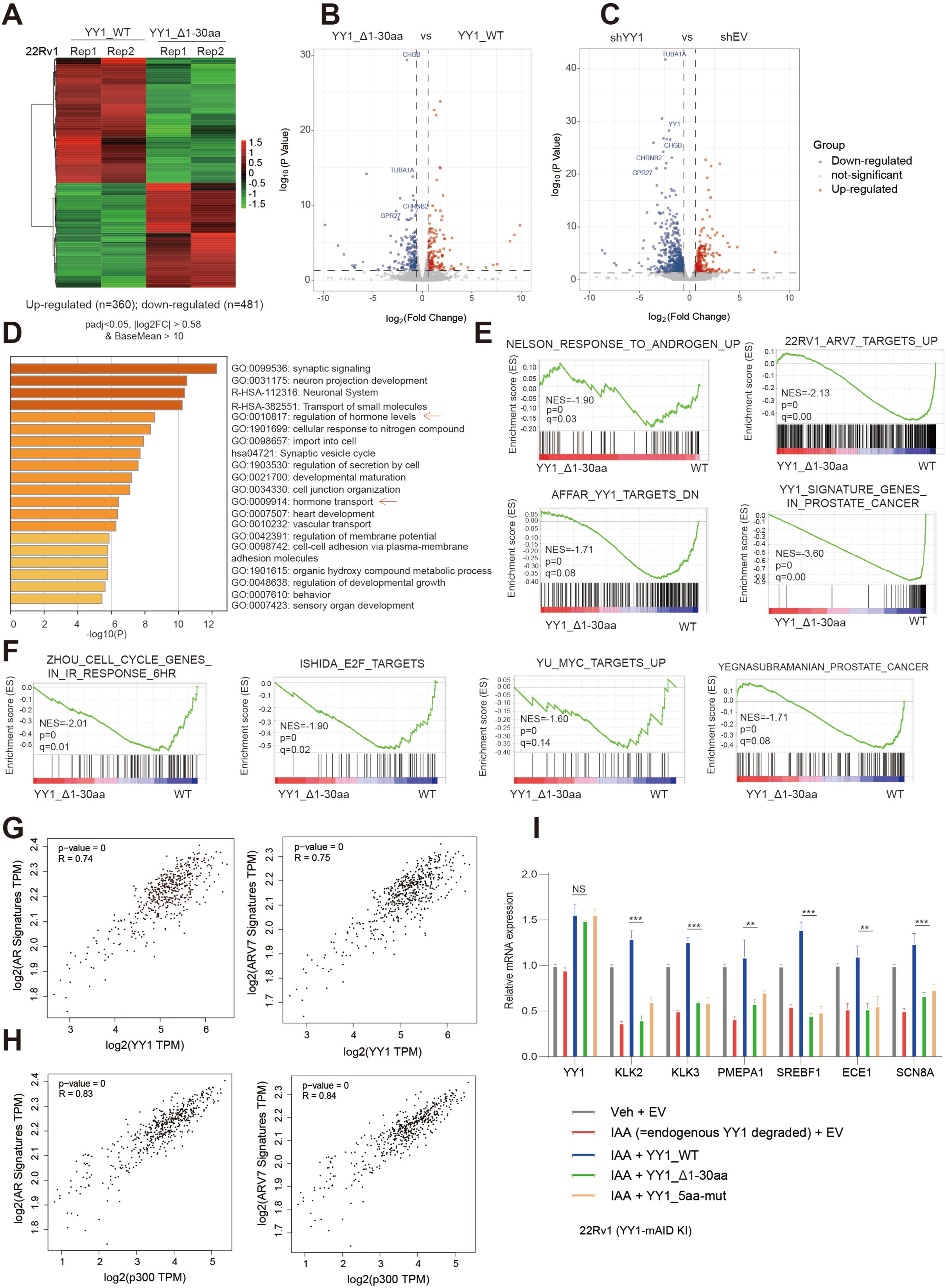
Transcriptomic profiling demonstrates a crucial involvement of YY1^AD^ for activating target oncogenes in CRPC cells. (A) Heatmap showing relative expression of the differentially expressed genes (DEGs) identified by RNA-seq in the YY1-degraded 22Rv1 cells (using the YY1-mAID KI cells, treated with 2.5 µM of 5-Ph-IAA for 48 hours to achieve YY1 degradation as shown in Figure 1E) that were rescued with WT YY1 versus YY1_Δ1-30aa (n = 2 two biological replicates per group). Threshold of DEG is set at the adjusted DESeq P value (padj) less than 0.01 and fold-change (FC) over 1.5 for transcripts with mean tag counts of at least 10. **(B-C)** Volcano plots showing transcriptomic alterations in the YY1-degraded 22Rv1 cells rescue by YY1_Δ1-30aa compared to WT YY1 (**B**), or in parental 22Rv1 cells with YY1 knockdown (KD) versus mock controls (EV; **C**). The most significantly altered transcripts are labeled and highlighted in the plots. (D) Gene Ontology (GO) analysis of 481 genes downregulated in 22Rv1 cells due to YY1_Δ1_30aa mutation relative to WT YY1 controls, highlighting pathways associated with AR signaling and cell proliferation. **(E-F)** Gene Set Enrichment Analysis (GSEA) using the RNA-seq profiles of YY1-degraded 22Rv1 cells shows that rescue with YY1-Δ1_30aa versus WT YY1 is positively correlated with the downregulation of androgen- or AR-V7-upregulated transcripts (**E**: top panels), the YY1 signature genes (**E**: bottom panels), as well as various oncogenic pathway genes **(F)** related to the cell cycle progression, E2F, c-MYC, and prostate tumorigenesis. **(G-H)** Pearson correlation analysis between mRNA expression levels of YY1 (**G**) or p300 (**H**) and the AR signature genes (left panels) and AR-V7 signature genes (right panels) in the TCGA prostate cancer cohort. (**I**) RT-qPCR of YY1 and selected downstream target genes in the YY1-degraded 22Rv1 cells, transduced with HA-tagged YY1, either WT or YY1^AD^ mutant (Δ1_30aa or 5aa_mut). Y-axis, presented as mean ± SD, shows the relative transcription level after normalizing to an internal control and to mock-treated samples. **, *P* < 0.01; ***, *P* < 0.001.

### Genome-wide profiling substantiates a role for YY1^1-30aa^ in potentiating H3K27ac levels globally in CRPC

Next, we sought to dissect how the N-terminal YY1 sequence (YY1^1-30aa^) acts to modulate the p300 activation-associated H3K27ac in CRPC. Here, we used the YY1-depleted models, which included cells carrying the mAID2-based YY1 degradation system (both 22Rv1 and VCaP cells) or those carrying the YY1-targeting shRNA (VCaP), for rescue with HA-tagged YY1, either WT or truncated YY1^Δ1–30aa^. Next, we performed the spike-in-controlled CUT&Tag assays for YY1 (with tag-specific antibody), H3K27ac and H3K27me3. Genome-wide correlational plots showed the bindings of WT YY1 and YY1^Δ1–30aa^ to be highly correlated in 22Rv1 and VCaP cells (**Fig. 5A; Supplementary Fig. S5A**). Both YY1 forms (WT and YY1^Δ1–30aa^) are widely and similarly distributed among the cis-regulatory elements such as promoters and putative enhancers (as annotated in **Supplementary Fig. S5B-C**). In contrast, we observed a significant reduction in genome-wide H3K27ac in both 22Rv1 (**Fig. 5B**) and VCaP (**Fig. 5C**) cells carrying the truncated YY1^Δ1–30aa^ relative to WT controls, resembling what was seen with YY1 loss (**Fig. 5B**, IAA vs. Veh; **Fig. 5C**, shYY1 vs. shEV). Meanwhile, overall H3K27me3 patterning remained largely unchanged in these cells (**Fig. 5D**). The H3K27ac changes were significant at the global level, as assessed by using reads densities at the called peaks (after normalization to respective spike-in controls; **Fig. 5E** and **Supplementary Fig. S5D**) and obvious at the classic AR target genes such as KLK2, KLK3 and PMEPA1 (**Fig 5F** and **Supplementary Fig. S5E-G**). Thus, YY1^1-30aa^ is dispensable for the YY1 genomic binding, which agrees with a notion that the C-terminal zinc fingers of YY1 mediate DNA binding (**Fig. 2G**); instead, our data show that YY1^1-30aa^ acts to promote H3K27ac at target sites.

**Figure 5.**
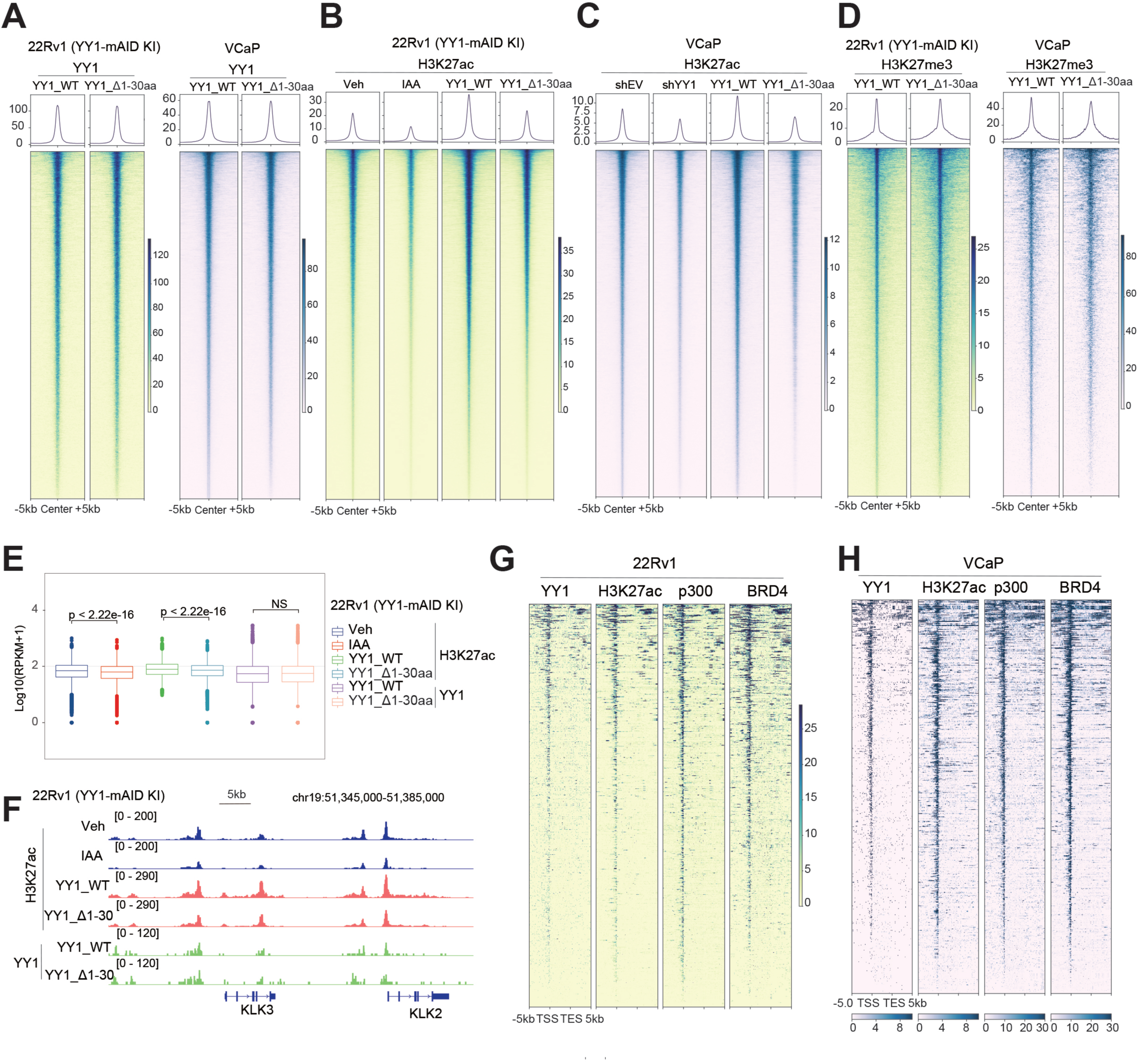
CUT&Tag profiling reveals an essential involvement of YY1^1-30aa^ for enhancing H3K27ac globally in CRPC. (A) Heatmap showing the anti-HA CUT&Tag signal densities (after spike-in control normalization), ±5 Kb around the called peaks in the YY1-depleted 22Rv1 (left) or VCaP (right) cells rescued with HA-tagged YY1, either WT or YY1_Δ1-30aa. **(B-C)** Heatmap showing the H3K27ac CUT&Tag signal densities (after spike-in control normalization), ±5 Kb around the called peaks in 22Rv1 **(B)** or VCaP cells **(C**), either mock-treated (column 1), YY1 depleted (column 2), or the YY1-depleted cells rescued with HA-tagged WT YY1 (column 3) or YY1_Δ1-30aa (column 4). YY1 depletion in the YY1-mAID 22Rv1 cells was achieved using the 5-Ph-IAA treatment while that in VCaP cells was done with shRNA-mediated YY1 KD. **(D)** Heatmap showing the H3K27me3 CUT&Tag signal densities (after spike-in control normalization), ±5 Kb around the called peaks in the YY1-depleted 22Rv1 (left) and VCaP (right) cells, rescued with HA-tagged WT YY1 or YY1_Δ1-30aa. **(E-F)** Plots for the overall H3K27ac and YY1 CUT&Tag signals (**E**; using the log2-transformed RPKM counts) and IGV views at the indicated gene locus **(F)** in 22Rv1 cells, either mock-treated, YY1 depleted, or the YY1-depleted cells rescued with HA-tagged WT YY1 or YY1_Δ1-30aa. **(G-H)** Heatmap showing overall binding of the indicated proteins at a suite of genes exhibiting the YY1_1-30aa dependency for their activation in 22Rv1 (**G**, n = 481) or VCaP cells (**H**, n = 403). TSS, transcriptional start site; TES, transcriptional end site.

Furthermore, our integrated studies using CUT&Tag and RNA-seq datasets showed that DEGs down-regulated due to YY1^Δ1–30aa^ in 22Rv1 (**Fig. 5G**; n = 481) and VCaP cells (**Fig. 5H**, n = 403) were generally co-bound by YY1, p300, H3K27ac and BRD4 (a histone acetylation reader and transcriptional coactivator). Concurrent with H3K27ac decrease at these genes upon YY1^Δ1–30aa^ in 22Rv1 and VCaP cells (**Fig. 5B**-**C**), there was an increase of H3K27me3 (**Supplementary Fig. S5H**). These observations suggest that the reduction of H3K27ac and an increase of H3K27me3 may collectively contribute to target gene down-regulation upon deletion of YY1^1-30aa^. Besides binding p300 as studied in this paper, YY1 was reported to interact with Polycomb Repressive Complexes as well^43–45^. Thus, the YY1^1-30aa^-mediated p300 interaction is likely to facilitate a functional switch of YY1 from the repressor to activator. In support, the downregulation of genes demarcated by H3K27me3 in prostate cancer is positively correlated with deletion of YY1^1-30aa^ relative to WT YY1 in CRPC cells (**Supplementary Fig. S5I-J**).

### YY1:p300 interaction acts to promote the AR/AR-V7-related oncogenic signaling in CRPC

AR-V7, a constitutively active AR form, activates the AR signaling in the absence of ligand and correlates with therapeutic resistance and poorer outcome seen with CRPC patients^46^. Have observing that YY1^1-30aa^ is crucial for activating the select classic AR/AR-V7 target genes (such as KLK2, KLK3 and PMEPA1; **Fig. 4I**), we next conducted a more comprehensive analysis of YY1- and AR/AR-V7-related cistrome in CRPC cells. Approximately 71% and 49% of AR-V7-bound sites were co-occupied by YY1 in 22Rv1 and VCaP cells respectively (**Fig. 6A**, **Supplementary Fig. S6A**), while about 38% and 5% of sites bound by ligand (DHT)-activated full-length AR exhibited the YY1 co-localization in these cells (**Fig. 6B**, **Supplementary Fig. S6B**). YY1 has been known as an AR cofactor^47^, but how it regulates AR-V7 hasn’t been explored before. An unbiased motif search analysis using the AR-V7-bound peaks in 22Rv1 cells discovered the YY1 motif among the top enriched motifs (**Supplementary Fig. S6C**). These results suggest a role of YY1 in regulating cistrome of the constitutively active AR-V7 in CRPC. As a strong support of this notion, we identified YY1 to be one of the enriched hits in the BioID-based AR-V7 interactome study in 22Rv1 cells (**Fig. 6C** and **Supplementary Fig. S6D**); additionally, co-IP detected the interaction between the exogenously expressed YY1 and AR-V7 in 293 cells (**Supplementary Fig. S6E**), as well as endogenously expressed YY1 and AR-V7 in CRPC cells (both 22Rv1 and VCaP; **Supplementary Fig. S6F**). To map out the YY1:AR-V7 interaction interface, we used various YY1 deletion constructs (**Supplementary Fig. S6G**) and found the region spanning amino acids 198–330 of YY1 to be crucial for AR-V7 interaction (**Fig. 6D)**. Genomic localization analysis showed that most of YY1- and AR-V7/AR-cobound peaks are localized at intergenic and intronic regions in CRPC cells (**Supplementary Fig. S6H-I**), indicating that YY1 mainly regulates the AR/AR-V7 signaling via enhancer activation. Indeed, peaks cobound by YY1 and AR-V7/AR were positive for H3K27ac, a marker for active enhancers, and p300 in 22Rv1 (**Fig. 6E-F)** and VCaP cells (**Supplementary Fig. S6J-K**).

**Figure 6.**
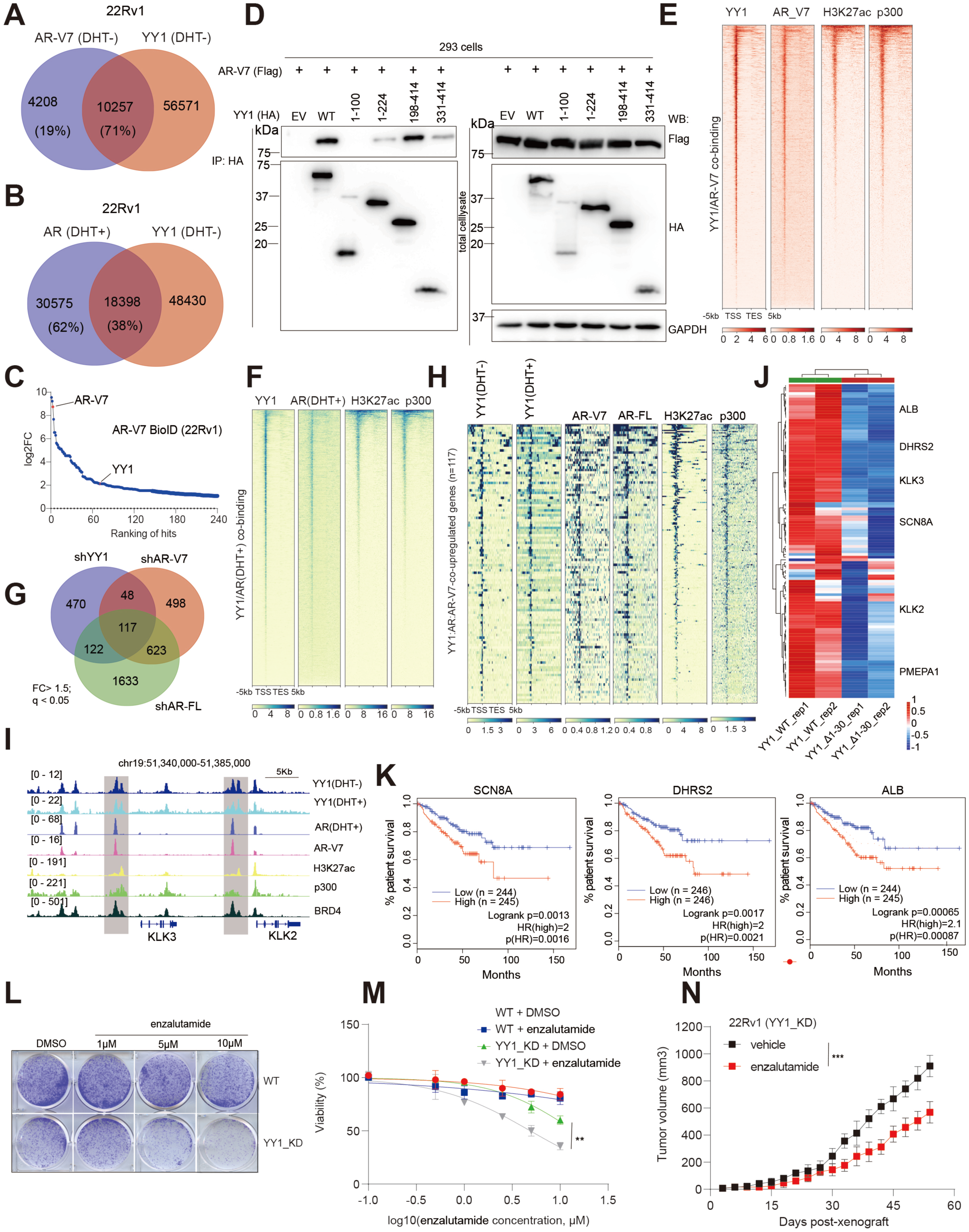
YY1:p300 interaction acts to promote the AR/AR-V7-related oncogenic signaling in CRPC, conferring therapeutic resistance. (A-B) Venn diagram showing the overlap between the called YY1 peaks and those of AR-V7 (**A**) or the ligand (DHT)-activated AR (**B**) peaks in 22Rv1 cells. **(C)** BioID identifies AR-V7-assoicated partner proteins in 22Rv1 cells, with the hits ranked by the FC of normalized spectral abundance factor (NSAF) relative to empty vector (EV) control. **(D)** Co-IP using the YY1 serial deletion constructs mapped out its AR-V7 interaction region to be an internal region (198-330aa). **(E-F)** Heatmap showing the YY1, H3K27ac, p300, and AR-V7 or AR binding signals, ± 5 kb from the centers of either YY1:AR-V7 (**E**) or YY1:AR (**F**) co-bound peaks identified in 22Rv1 cells. TSS, transcriptional start site; TES, transcriptional end site. **(G)** Venn diagram using DEGs, identified by RNA-seq to be down-regulated in 22Rv1 cells post-depletion of either YY1, AR, or AR-V7 (n = 2 biologically independent experiments). FC, fold change. **(H)** Heatmap showing overall binding of the indicated protein at YY1:AR:AR-V7-co-upregulated genes (n = 117; defined in **G**). **(I)** IGV views showing the enrichment of the indicated protein at *KLK3* and *KLK2* in 22Rv1 cells. **(J)** Heatmap showing relative expression of the 117 genes co-activated by YY1, AR and AR-V7 in the YY1-depleted 22Rv1 cells rescued with WT YY1 versus YY1_Δ1-30aa mutant. Sample information was indicated on the bottom of the heatmap (n = 2 replicate per group). **(K)** Kaplan–Meier disease-free survival analysis based on the expression of *SCN8A* (left), *DHRS2* (middle) or *ALB* (right) in patient samples from the TCGA cohort of prostate cancers. Statistical significance was determined by log-rank test and shown. **(L)** Colony formation by 22Rv1 cells, either mock (WT) or with YY1 depleted (YY1_KD), in the presence of the indicated enzalutamide concentration. **(M)** Plots showing the growth-inhibitory effect of various used concentration of enzalutamide (x-axis in log10-converted value; treated for 8 days) in 22Rv1 cells, either mock (WT) or with YY1 depleted (YY1_KD). The y-axis shows relative cell growth after normalization to DMSO-treated cells (n = 3 independent treatment experiments; presented as the mean ± SD). **, *P* < 0.01. **(N)** Growth kinetics of the xenografted tumors after subcutaneous transplantation of the YY1-depleted 22Rv1 cells (YY1-KD) into castrated NSG mice, treated with either vehicle (black) or enzalutamide (red; (n = 8 per group). ** *P* < 0.005, *** *P* < 0.001.

Next, we utilized the RNA-seq profiles of 22Rv1 cells before and after KD of each one of the oncogenic TFs studied herein (namely, YY1, AR and AR-V7) and identified a set of 117 genes that are co-activated by these TFs (**Fig. 6G**). Such YY1:AR:AR-V7 co-upregulated genes were generally bound directly by these oncogenic TFs (YY1 and AR/AR-V7), as well as H3K27ac and p300 (**Fig. 6H**), as exemplified by what was observed at KLK3, KLK2, PMEPA1 and SCN8A (**Fig. 6I** and **Supplementary Fig. S6L-M**). Transcriptional activation of these 117 genes was also dependent on YY1’s p300-interacting domain (namely, YY1^1-30aa^), as shown by comparison of RNA-seq profiles of cells with WT YY1 relative to the truncated mutant YY1^Δ1-30aa^ (**Fig. 6J**). Such YY1:AR:AR-V7 co-upregulated signature genes were clinically relevant. For example, high expression of some of these genes (such as SCN8A, DHRS2 and albumin) was correlated with poor clinical outcomes of prostate cancer patients (**Fig. 6K**).

Given that YY1 potentiates the AR-V7 signaling, a clinically relevant pathway leading to therapeutic resistance seen in CRPC, we further asked if the loss of YY1 would overcome treatment resistance. Towards this end, we treated parental and 22Rv1 cells with YY1 KD *in vitro* with an increasing dose of enzalutamide, a mainstream therapy in the clinic for CRPC, and found that 22Rv1 cells with the YY1 KD become sensitive to enzalutamide treatment compared with WT controls (**Fig. 6L-M**). To further assess if the YY1 depletion may resensitize CRPC to the enzalutamide treatment *in vivo*, we inoculated the YY1-depleted 22Rv1 cells in the NOD/scid/gamma (NSG) mice, and once the tumors grew out (reaching approximately 100 mm³ in averaged sizes), we initiated the enzalutamide treatment. When compared with vehicle-treated controls, the enzalutamide treatment significantly inhibited growth of the YY1-depleted tumor xenografts (**Fig. 6N**).

Taken together, our results demonstrate that YY1 acts as a critical partner of AR/AR-V7 and activates the AR/AR-V7 signaling in CRPC, which at least partly involves YY1^1-30aa^ to release p300 auto-inhibition and promote H3K27ac at the AR/AR-V7-bound genomic sites. The loss of YY1 sensitizes CRPC cells to enzalutamide, indicative of a potential strategy to overcome therapeutic resistance caused by the constitutively active AR variant in CRPC patients.

### The YY1^1-30aa^-mediated p300 activation is critically involved in prostate tumorigenesis *in vitro* and *in vivo*

Lastly, we sought to define the role of YY1’s p300-interacting domain (YY1^1-30aa^) in prostate tumorigenesis. We first conducted rescue experiment in the YY1-KD 22Rv1 cells with either WT YY1 or YY1^Δ1-30aa^ and found that the latter truncated mutant failed to rescue tumor cell proliferation (**Fig. 7A-B**) and colony formation in soft agar, a surrogate assay of attachment-independent growth (**Fig. 7C** and **Supplementary Fig. S7A**). Relative to WT YY1, YY1^Δ1-30aa^ also failed to rescue the phenotypes caused by YY1 depletion, including the cell cycle progression defects, notably, the increase in G1-S phase cells and decrease of G2-M phase cells (**Fig. 7D**), as well as elevated apoptosis as assessed by FACS (**Fig. 7E)** and measurement of cleaved caspases and PARP levels (**Supplementary Fig. S7B**). Such failure to rescue the phenotypes caused by YY1 depletion were also observed in VCaP cells carrying YY1^Δ1-30aa^, when compared to WT controls (**Supplementary Fig. S7C-I**). We further examined the requirement of YY1^1-30aa^ for CRPC tumor growth *in vivo*. Here, we established the subcutaneous tumor xenografts in the castrated NSG mice with 22Rv1 cells, which were expressed with an inducible YY1 shRNA to deplete endogenous YY1 followed by restoration of YY1, either WT or YY1^Δ1-30aa^. The administration of doxycycline efficiently induced YY1 depletion *in vivo* (**Supplementary Fig. S7J: top panel**) and the bioluminescence imaging of live mice showed that YY1 depletion suppressed tumor growth *in vivo* as well (**Fig. 7F-H**; shEV vs. shYY1/KD). Meanwhile, WT YY1, but not its truncated form (YY1^Δ1-30aa^), restored growth of the 22Rv1 xenografted tumors (**Fig. 7F-H**; YY1_WT versus YY1_Δ1-30aa). Furthermore, the apoptosis marker cleaved-caspase3 was significantly increased in the isolated tumor samples with YY1 KD or YY1^Δ1-30aa^, when compared with the respective controls (**Fig. 7I-middle**). In contrast, KLK3/PSA, a classic prostate cancer biomarker, showed the opposite trend and is positively correlated with YY1 functionality, as assessed by IHC staining (**Fig. 7I-right**) and WB of isolated tumor samples (**Supplementary Fig. S7J**, see panel of KLK3). In consistent with *in vitro* results, YY1 depletion and rescue with YY1^Δ1-30aa^ also led to global decrease of H3K27ac in tumors in vivo, when compared with rescue with WT controls (**Supplementary Fig. S7J**, see panel of H3K27ac).

**Figure 7.**
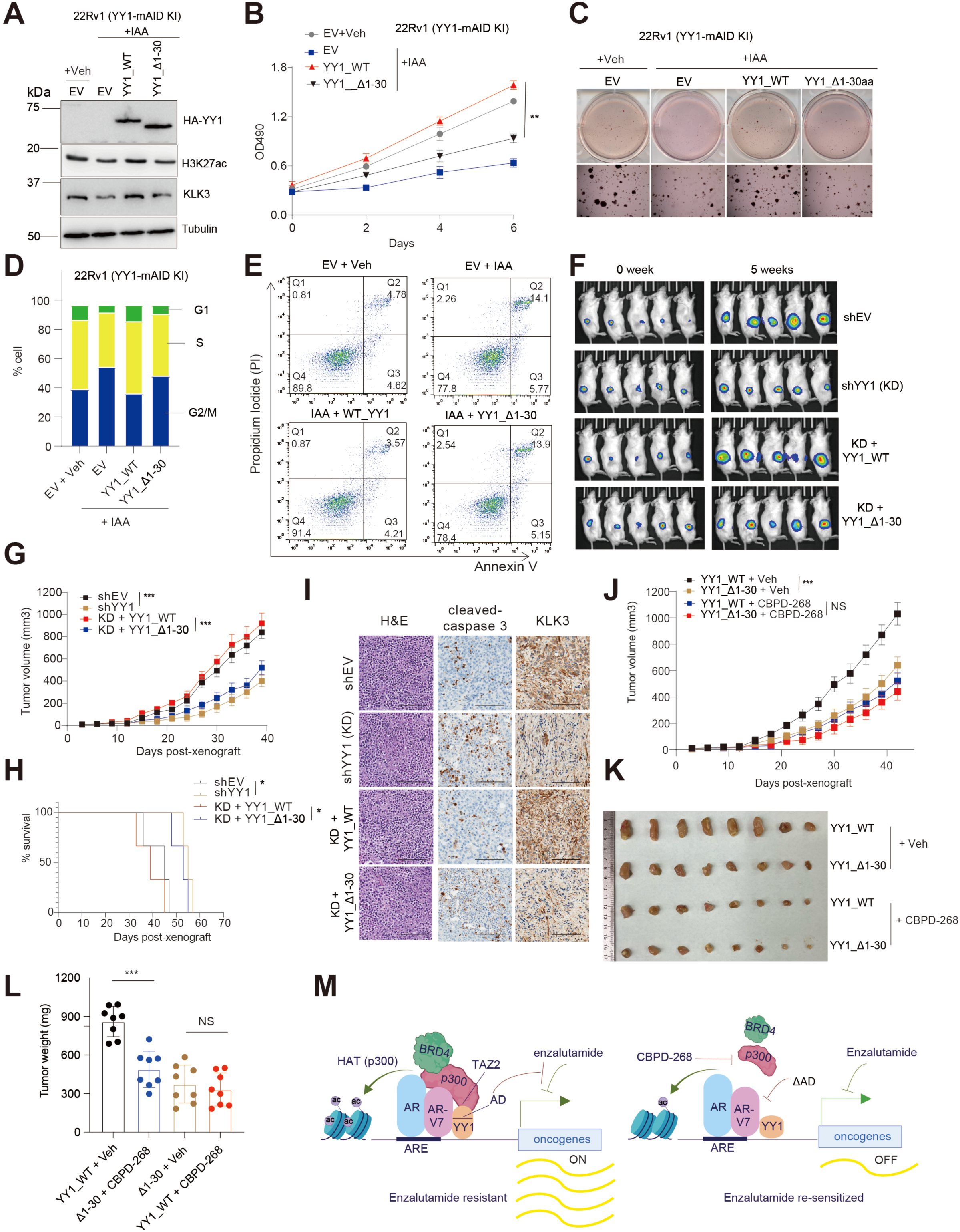
The YY1^1-30aa^-mediated p300 activation is critically involved in prostate tumorigenesis *in vitro* and *in vivo*. (A–E) WB for the indicated protein (**A**), measurement of cell proliferation (**B**) and soft agar-based growth (**C**), cell cycle analysis (**D**), as well as representative Annexin-V and Propidium Iodide (PI) staining (**E**) using 22Rv1 cells with the KI of YY1-mAID, treated with 5-Ph-IAA for YY1 depletion and then rescued with either vector mock (EV), WT YY1, and YY1_Δ1-30aa, compared to mock treatment (veh/EV); n = 3 independent treatment experiments (\*\**P* < 0.01). Q1, Q2, Q3 and Q4 in **panel E** represent cells undergoing necrosis, late-stage apoptosis and early-stage apoptosis and live cells, respectively. **(F-I)** Representative bioluminescence images of live animal (**F**) showing the 22Rv1 cell-xenografted tumors in NSG mice (n = 8 per group) before (0 week) and 5 weeks post-depletion of YY1 *in vivo* (doxycycline induced), quantification of tumor size (**G**), Kaplan– Meier curve of tumor-bearing animals (**H**), as well as Hematoxylin and Eosin (H&E) and immunohistochemistry (IHC) staining of the indicated protein (cleavaged caspase 3 and KLK3) in the xenografted tumors collected from NSG mice (**I;** scale bar, 25 μm). Castrated NSG mice were subject to subcutaneous transplantation of each one of four different luciferase-expressing 22Rv1 cells, which were stably transduced with either empty vector control (shEV) or a YY1-targeting shRNA (doxycycline-inducible shYY1 for YY1_KD), or the latter shYY1-bearing cells with pre-rescue of either WT YY1 (KD + YY1_WT) or Δ1-30aa (KD + YY1_ Δ1-30). Log rank test was used for **H** (\**P* < 0.05; *** *P* < 0.001). (**J-L**) Growth kinetics (**J**) as well as representative images (**K;** at the end of the study) and weight (**L;** at the end of the study) of tumors post-xenografting of 22Rv1 cells (expressing YY1_WT or YY1_Δ1-30 in the YY1-depleted background as in panels **F-I**) into castrated NSG mice, treated with vehicle (Veh) or CBPD-268 (n = 8 per group; \*\*\**P* < 0.001). CBPD-268 was administered orally at a dose of 1 mg/kg per day for five days each week (Monday to Friday), as detailed in Methods. **(M)** Left: Schematic model illustrating a regulatory axis involving interaction of YY1^AD^:p300^TAZ2^:AR-V7 in advanced prostate cancers, which acts to enhance p300 activity *in cis* to modulate the landscape of histone acetylation and robust transcription of oncogenes downstream of YY1 and/or AR; this axis also activates AR signaling to confer malignant growth and enzalutamide resistance seen in CRPC. Left: Targeting YY1^AD^ or p300 disrupts this above axis, reduces histone acetylation, and restores sensitivity to enzalutamide.

To explore whether the in vivo growth difference between xenografted tumors expressing the YY1^Δ1-30aa^ mutant relative to WT YY1 is dependent on p300 activity, we employed the recently disclosed degrader of p300/CBP, CBPD-268^47^. Towards this end, we established subcutaneous tumor xenografts with 22Rv1 cells expressing WT YY1 or YY1^Δ1-30aa^ as above in NSG mice. When averaged tumor sizes reached approximately 100mm³, the mice were treated orally with the vehicle control or CBPD-268 (at a dose of 1mg/kg for 5 days per week). Here, the YY1^Δ1-30aa^ mutant relative to WT resulted in significant tumor growth inhibition in the vehicle-treated groups; meanwhile, no significant difference in tumor growth was observed between the WT YY1 and YY1^Δ1-30aa^ groups receiving the CBPD-268 treatment (**Fig. 7J–L**). These results point to a phenotypic resemblance in the CBPD-268-treated YY1_WT group and the untreated YY1^Δ1-30aa^ group (**Fig. 7J–L**), supporting the effect of YY1^1-30aa^ is through the p300 interaction.

Taken together, our results strongly support that the p300-interacting YY1^1-30aa^ is critical for CRPC cell proliferation in vitro and xenografted CRPC tumor growth in vivo.

## Discussion

Genetic and epigenetic abnormalities, caused often by mis-regulation and mutation of DNA-binding factors and/or chromatin modulators, play central roles during the initial pathogenesis and advanced progression of aggressive human cancers such as lethal CRPC. It is known that the AR signaling hyper-activation (notably, via acquisition of AR-V7, amplification of AR, amongst others)^2,5–10^, as well as over-expression of YY1^12^, is common in CRPC patients^12,13,15^, underscoring an urgent need for alternative approaches to block these oncogenic pathways. Likewise, over-expression and/or activation of p300 (a histone acetyltransferase)^13,15^ and EZH2 (a histone methyltransferase^48,49^), as well as loss-of-function mutation of SWI/SNF (a chromatin-remodeling complex)^50^, were recurrently detected in clinic cohorts of advanced prostate cancers. Dissecting how exactly these genetic and epigenetic aberrations synergize in establishing a malignant tumor state shall not only improve current understanding of the biology underlying hard-to-treat cancers but, importantly, may suggest the attractive anti-cancer strategies.

In this study, we report that **<u>(i)</u>** in CRPC, YY1, but not other tested prominent oncogenic TFs (such as AR and related AR-V7, or a pioneering factor FOXA1), enhances the p300-mediated histone acetylation globally; **<u>(ii)</u>** based on the AlphaFold2-based structural modeling and subsequent biochemical assays, we show that, while YY1’s AD (YY1^AD^, which is evolutionarily conserved) is dispensable for p300’s chromatin binding, it is essential for directly binding p300’s TAZ2 domain to relieve p300 autoinhibition, thereby promoting p300’s HAT activity *in cis* (assessed by p300 trans-autoacetylation) and H3K27 hyperacetylation for gene activation (assessed by *in vitro* assays using the purified recombinant protein and luciferase-based assays in cells). **<u>(iii)</u>** Our integrative genomic studies substantiated the requirement of the YY1^AD^:p300^TAZ2^-mediated signaling for enhancing histone acetylation at target chromatin, for promoting robust transcription of downstream oncogenes, and for sustaining aggressive growth and drug resistance seen with CRPC models (see a model in **Fig. 7M-left panel**); **<u>(iv)</u>** critically, we have discovered YY1 as a functional partner of AR/AR-V7 in CRPC, contributing to the AR signaling hyperactivation and transcriptional programs that confer the resistance to androgen-targeted therapies (enzalutamide, a clinical anti-androgen drug). This notion is supported by our unbiased motif search analysis using TF-binding peaks in the genome, mass spectrometry-based TF interactome study and co-IP verification, as well as in-depth TF cistrome analyses that not only unveiled the substantial co-occupancies of YY1 and AR/AR-V7 at their genomic binding sites (assessed by CUT&Tag) but also defined a suite of direct target genes co-upregulated by YY1, AR/AR-V7 and p300 (assessed by integrated RNA-seq and CUT&Tag in **Fig. 6**). We found a protein region spanning the amino acids 198–330 of YY1 to be involved in the interaction with AR-V7, which merits further investigation in future. **<u>(v)</u>** In the clinic, a majority of patients receiving the androgen-targeted therapies (enzalutamide) ultimately progress to fatal stages (i.e., CRPC), despite mild extension of their survival. Here, we show that loss of YY1^AD^ led to the suppressed CRPC growth *in vitro* and *in vivo* in a p300-dependent fashion, and such a functional blockade of YY1 also significantly resensitized CRPC to the enzalutamide treatment (see a model in **Fig. 7M-right panel**). Additionally, the YY1-overexpressed CRPC cells were more sensitive to p300 blockade *in vivo* (**Fig. 7J-K**), indicative of another alternative anti-tumor strategy.

In summary, our observations highlight a crucial involvement of the crosstalk between cancer-associated TFs and chromatin factors for generating more aggressive phenotypes seen in CRPC. YY1, when overexpressed in advanced prostate cancers, functions to relieve p300 autoinhibition *in cis* to modulate the global landscape of histone acetylation and enhance the oncogenic signaling downstream of YY1 and AR/AR-V7, which hasn’t reported before. CRPC remains an unresolved clinical challenge. Based on our findings, further investigation is warranted to develop a means to block YY1^AD^:p300^TAZ2^ interaction as a strategy for suppressing this critical oncogenic signaling in CRPC. Also, p300 inhibition has recently emerges as a compelling strategy for treating advanced prostate cancers^15^ — CCS1477 is a p300/CBP bromodomain inhibitor currently under the clinical evaluation^15^, while two orally bioavailable PROTAC degraders against p300/CBP have been recently developed^47,51^, including CBPD-268 that we evaluated herein with *in vivo* dosing study. We are optimistic of the above therapeutic strategies indicated in this study, which shall assist in development of the new treatment for the otherwise drug-resistant CRPC patients.

