## Supplemental figures and legends for "YY1 relieves p300 autoinhibition to promote histone acetylation in advanced prostate cancer, enhancing the oncogenic signaling of Androgen Receptor Splice Variant 7"

Fig S1 Xu et al

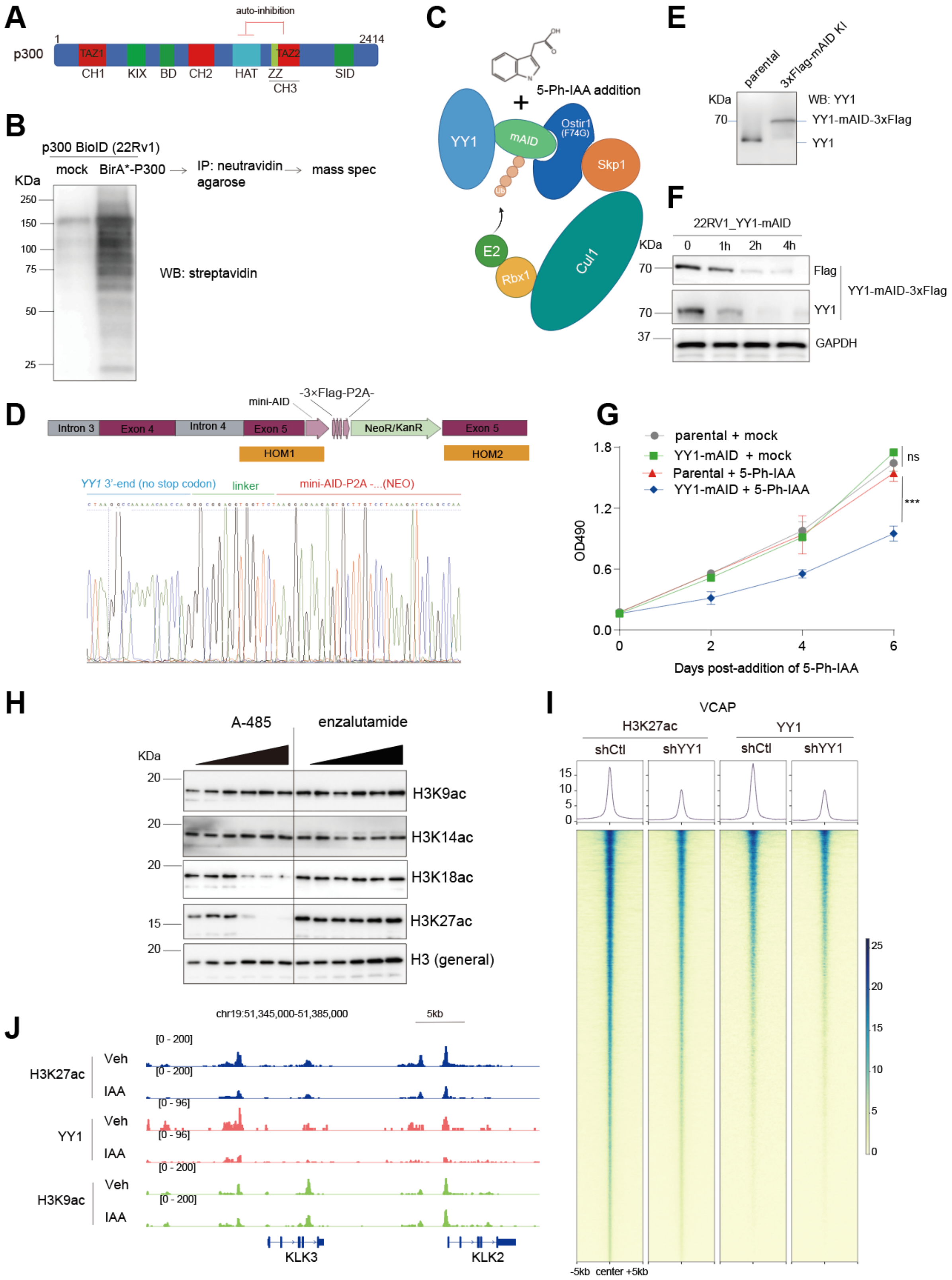


**Supplementary Fig. 1. YY1, and not other tested oncogenic TFs (AR, AR-V7 or FOXA1), enhances the p300-mediated histone acetylation globally in CRPC.**

**(A)** Schematic of the p300 functional domains. TAZ2, a TF-interacting domain in p300, contributes to an auto-inhibitory mechanism by negatively regulating the enzymatic activity of the histone acetyltransferase (HAT) domain[1].

**(B)** WB of the biotinylated protein species using a streptavidin-HRP conjugate antibody in 22Rv1 cells, which were stably transduced with a biotin ligase (BirA*) only (mock controls; lane 1) or BirA*-p300 fusion (lane 2), followed by treatment with 50 μM of biotin for 24h.

**(C)** Schematic illustration depicting the mini AID2 system (mAID) for targeted degradation of endogenous YY1 protein, induced by addition of 5-Ph-IAA.

**(D)** Top: Schematic showing the pFETCh-based genomic targeting strategy employed to insert a 3×Flag-mAID-P2A-NeoR$^+$ cassette in-frame to the C-terminus of the YY1 gene through homologous recombination using the two homology arms (HOM1 and HOM2).

Bottom: Representative Sanger sequencing traces confirmed precise recombination and successful knockin (KI) of the 3×Flag-mAID-P2A-NeoR$^+$ cassette into the YY1 loci of 22Rv1 cells, creating a stable line harboring homozygous KI allele of YY1-3×Flag-mAID.

**(E)** WB using anti-YY1 antibody in 22Rv1 cells, either parental or those with the YY1-3×Flag-mAID KI alleles, detecting normal YY1 and YY1-3×Flag-mAID fusion in correct sizes.

**(F)** WB for the indicated protein using total lysates from 22Rv1 cells carrying the YY1-3×Flag-mAID KI alleles, treated with 5 uM of 5-Ph-IAA for 0, 2 and 4 hours.

**(G)** Measurement of proliferation of 22Rv1 cells, either parental or those carrying the KI of YY1-3×Flag-mAID, treated with vehicle (veh) or 5 uM of 5-Ph-IAA for the indicated duration (n= 3 replicates per group).

**(H)** A-485, but not enzalutamide, decreased global levels of H3K27ac and H3K18ac in 22Rv1 cells. Cells were treated after 48 hours with 5-fold dilutions of A-485 or enzalutamide, starting at 10 μM. DMSO served as a control.

**(I)** Heatmap of the H3K27ac and YY1 CUT&Tag read densities at the called H3K27ac peaks in VCaP cells, either with YY1 knockdown (KD; sh#94) or mock treated (EV).

**(J)** Integrative Genomics Viewer (IGV) views of YY1, H3K27ac and H3K9ac at KLK3 and KLK2 in 22Rv1 cells, treated with 5-Ph-IAA in comparison with Vehicle (Veh).

**Fig S2 Xu et al**

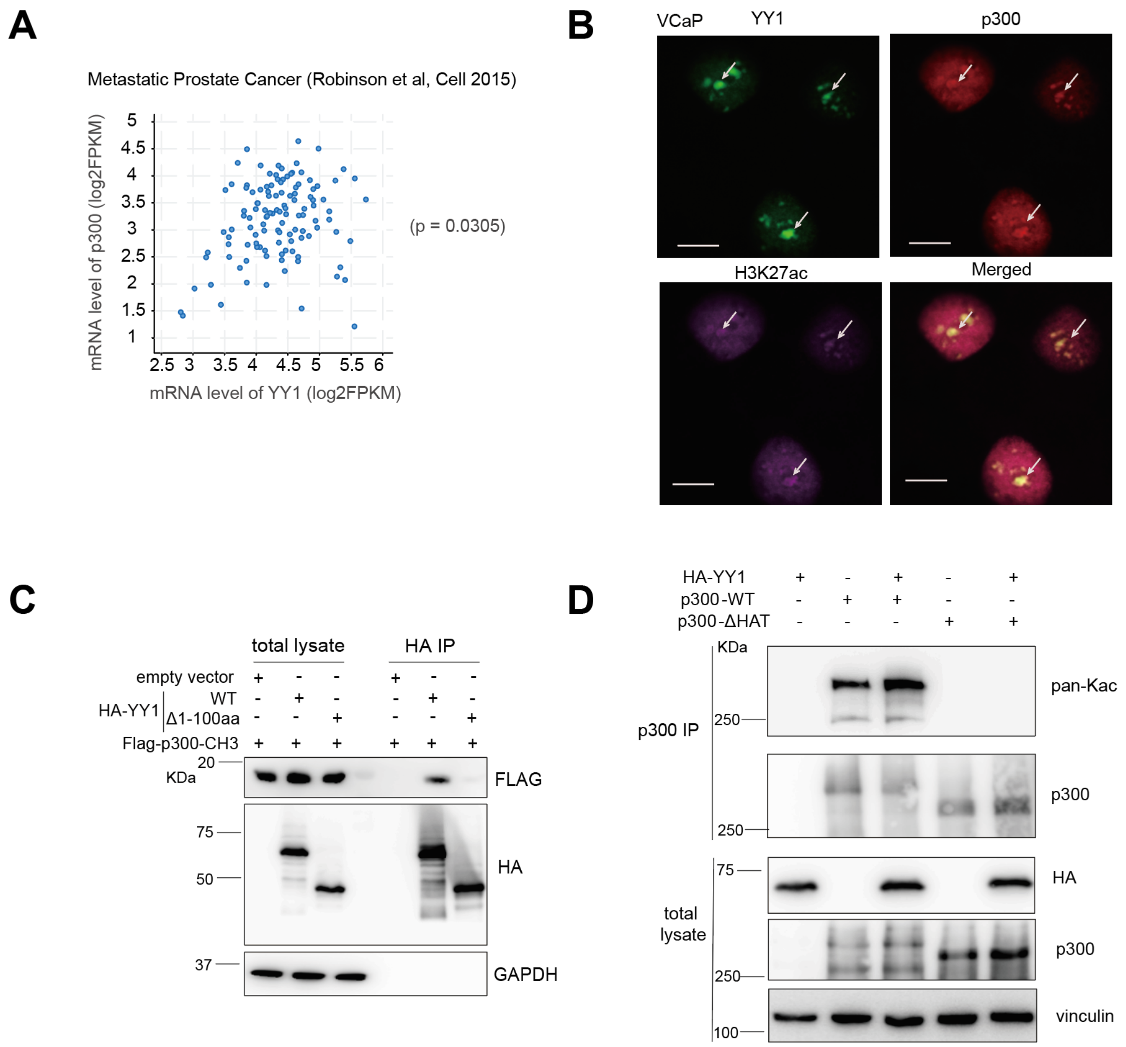


**Supplementary Fig. 2. YY1's N-terminus interacts with p300's CH3 module, relieving p300 autoinhibition to promote p300 trans-autoacetylation and substrate acetylation for gene activation.**

**(A)** Pearson correlation plot using the mRNA expression levels of YY1 and p300 in the metastatic prostate cancer database[2] of Robinson et al (2015).

**(B)** Immunofluorescence (IF) of the indicated protein in VCaP cells. Representative results from three independent experiments. Arrows highlight the co-localization. Scale bar, 10 μm.

**(C)** CoIP for interaction between exogenously expressed Flag-tagged p300$^{CH3}$ and HA-tagged YY1, either WT or a N-terminal truncated mutant (Δ1-100aa) in 293 cells.

**(D)** Assessing p300 trans-autoacetylation levels (based on WB with a pan-Kac antibody) post-IP of p300 from 293 cells co-transfected with tagged p300, either WT or the internal HAT domain-deleted form **(**p300_ΔHAT), and HA-tagged WT YY1. Both IP samples (top panels) and whole-cell lysates (bottom panels) were collected and used for WB of the indicated protein such as p300 and HA-YY1.

Fig S3 Xu et al

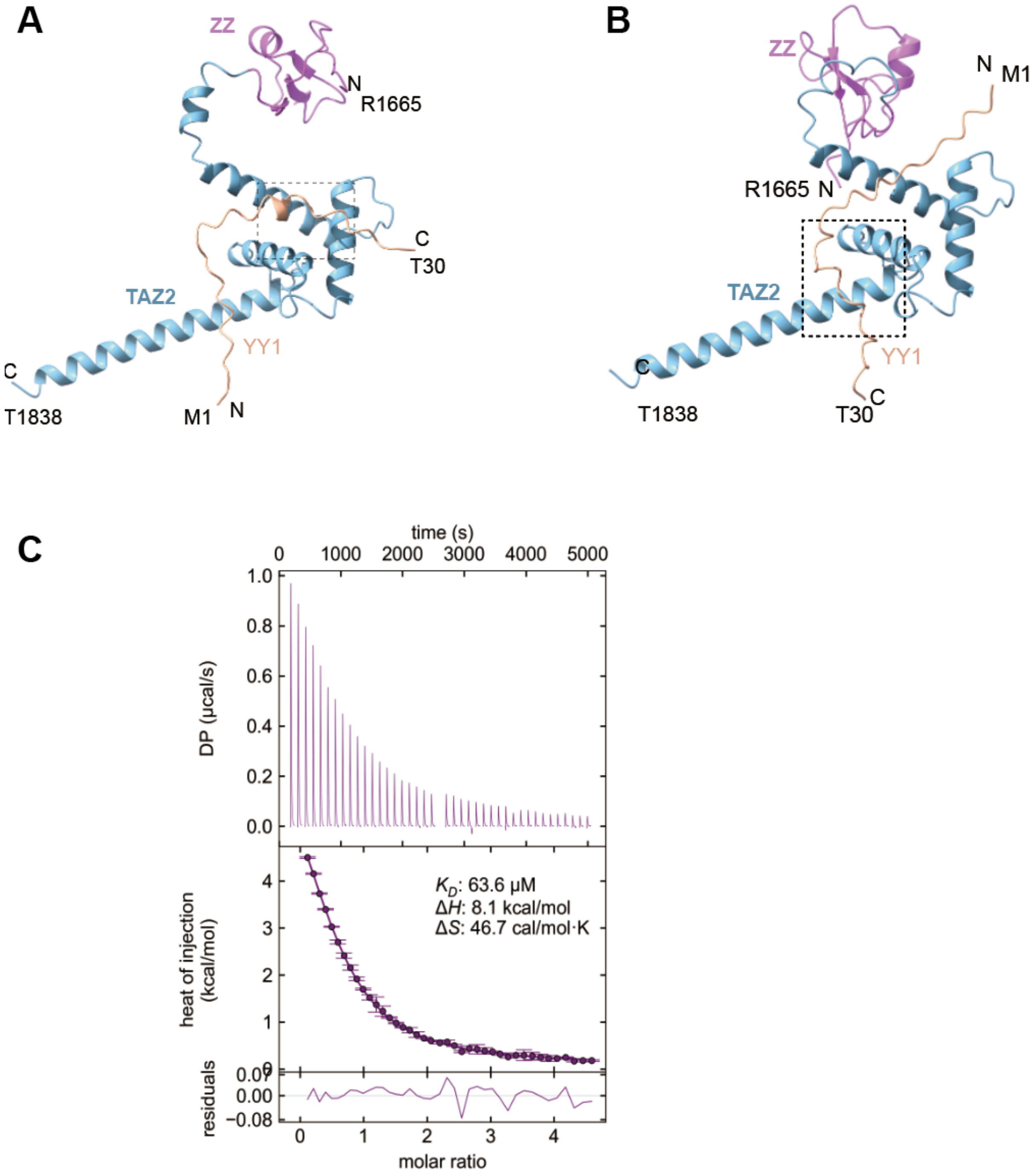


**Supplementary Fig. 3. Structural and biochemical analyses of the p300$^{CH3}$:YY1$^{AD}$ interaction.**

**(A-B)** AlphaFold2 predicting the two highly confident structural models of the p300$^{CH3}$ and YY1$^{1-30}$ complex, i.e., Model 1 (**A**) and Model 2 (**B**).

**(C)** Isothermal Titration Calorimetry (ITC) binding curves measuring the binding affinity between the purified recombinant protein of YY1$^{1-30}$ and p300$^{TAZ2}$.

Fig S4 Xu et al

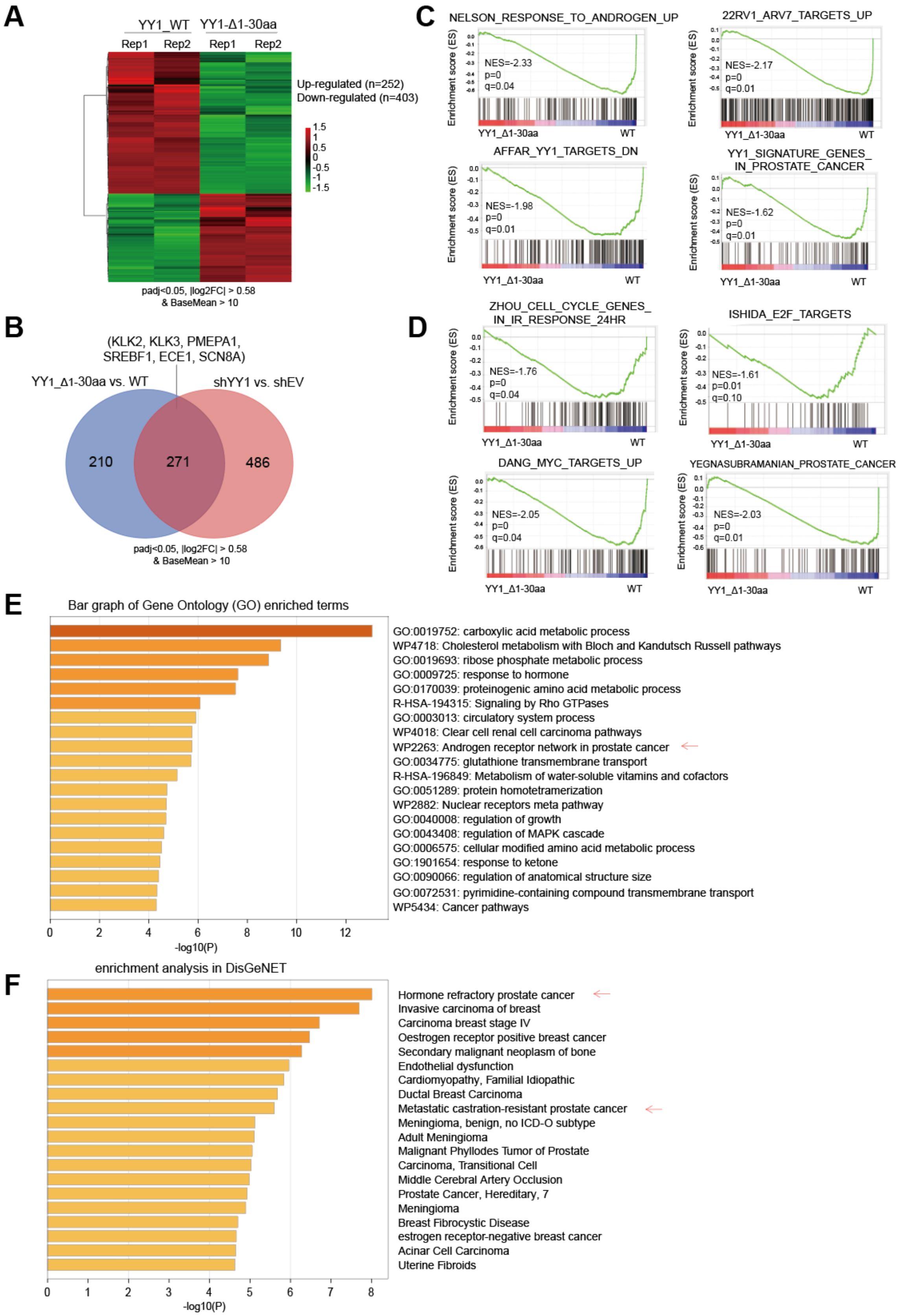


**Supplementary Fig. 4. Transcriptomic profiling demonstrates a crucial involvement of YY1$^{AD}$ for activating target oncogenes in CRPC cells.**

**(A)** Heatmap showing relative expression of the differentially expressed genes (DEGs) identified by RNA-seq in VCaP cells after the knockdown (KD) of YY1 and then rescue with YY1_Δ1-30aa, relative to WT YY1 controls (n = 2 two biological replicates per group). Threshold of DEG is set at the adjusted DESeq P value (padj) less than 0.01 and fold-change (FC) over 1.5 for transcripts with mean tag counts of at least 10.
**(B)** Venn diagram showing the overlap between downregulated DEGs in 22Rv1 cells due to the YY1_Δ1-30aa mutation (n = 481, left; YY1_Δ1-30aa vs. WT) and those due to YY1 KD (n = 757, right; shYY1 vs. shEV), compared with their respective controls as defined by RNA-seq. Threshold of DEG is set at the adjusted DESeq P value (Padj) less than 0.05 and FC over 1.5 for transcripts with mean tag counts of at least 10.
**(C)** Geneset enrichment analysis (GSEA) demonstrating that, in VCaP cells, the YY1_Δ1-30aa mutation relative to WT controls is positively associated with the downregulation of genes linked to the androgen- or AR-V7-upregulated genes (top panels), as well as the YY1-related signature genes (bottom panels).
**(D)** GSEA showing that, in VCaP cells, the YY1_Δ1-30aa mutation relative to WT controls is positively associated with the downregulation of gene pathways associated with the cell cycle progression, E2F signaling, MYC, and prostate cancer development.
**(E-F)** Gene ontology (GO; **E**) and enrichment of the DisGeNet category (**F**) analyses by using the downregulated DEGs (n = 403) in the YY1-depleted in VCaP cells rescued with YY1_Δ1-30aa, relative to WT YY1.

Fig S5 Xu et al

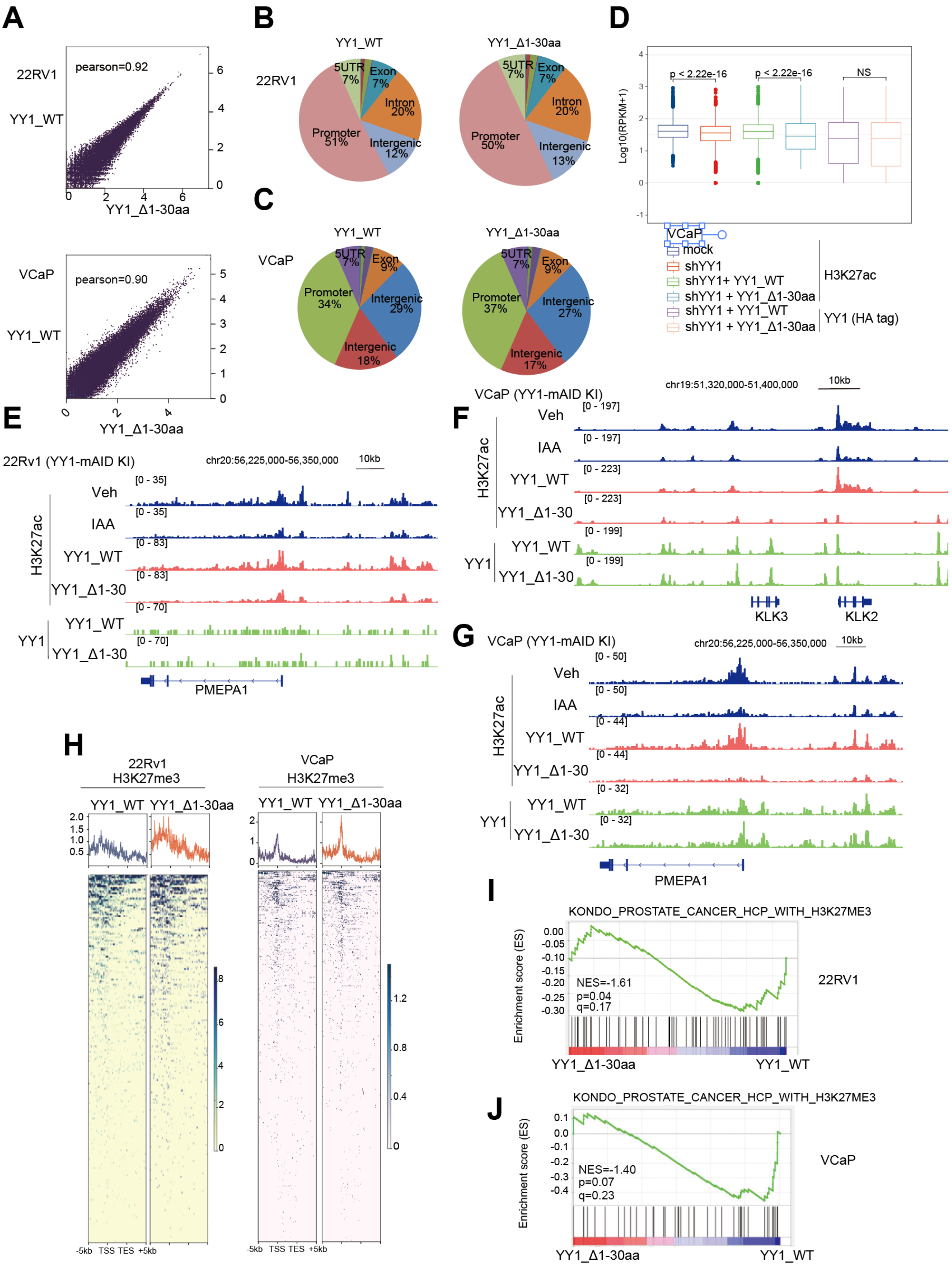


**Supplementary Fig. 5. CUT&Tag profiling reveals an essential involvement of YY1$^{1\text{-}30aa}$ for enhancing H3K27ac globally in CRPC.**

**(A)** Pearson correlation analysis of the WT YY1 and YY1_Δ1-30aa CUT&Tag signals in 22Rv1 (top) and VCaP (bottom) cells.
**(B-C)** Pie chart showing genomic annotation of the WT YY1 (left) and YY1_Δ1-30aa (right) CUT&Tag peaks in 22Rv1(**B**) and VCaP (**C**) cells.
**(D)** Plotting using the log2-transformed RPKM counts of the H3K27ac and anti-HA-YY1 CUT&Tag signals in VCaP cells. The used VCaP cells were stably transduced with control vector (mock) or YY1-targeting shRNA (shYY1 for YY1 depletion), or the latter YY1-depleted cells pre-arescued with HA-tagged YY1, either WT (shYY1 + YY1_WT) or the Δ1-30aa mutant (shYY1 + YY1_ Δ1-30aa).
**(E-G)** IGV views of CUT&Tag profiles for H3K27ac and YY1 (anti-HA) at the indicated gene locus in 22Rv1 **(E)** and VCaP cells (**F-G**). The used cells carried the engineered YY1-mAID knock-in (KI) alleles and were treated with either mock (Veh) or 5-Ph-IAA (IAA; to induce YY1 degradation), or the YY1-degraded cells pre-rescued with HA-tagged YY1, either WT (IAA + YY1_WT) or Δ1-30aa mutant (IAA + YY1_Δ1-30aa).
**(H)** Heatmap showing the H3K27me3 densities around the genes found to be downregulated due to the YY1_Δ1-30aa mutation relative to WT controls in 22Rv1 (left, n = 481) or VCaP cells (right, n = 403). The overall H3K27me3 increase in cells expressing the mutant versus WT YY1 indicated that a repressive chromatin environment underlies the target gene downregulation.
**(I-J)** GSEA showing that the YY1_Δ1-30aa mutation relative to WT controls is correlated with the downregulation of H3K27me3-demarcated genes in 22Rv1 **(I)** and VCaP **(J)** cells, suggesting an epigenetic repression mechanism.

Fig S6 Xu et al

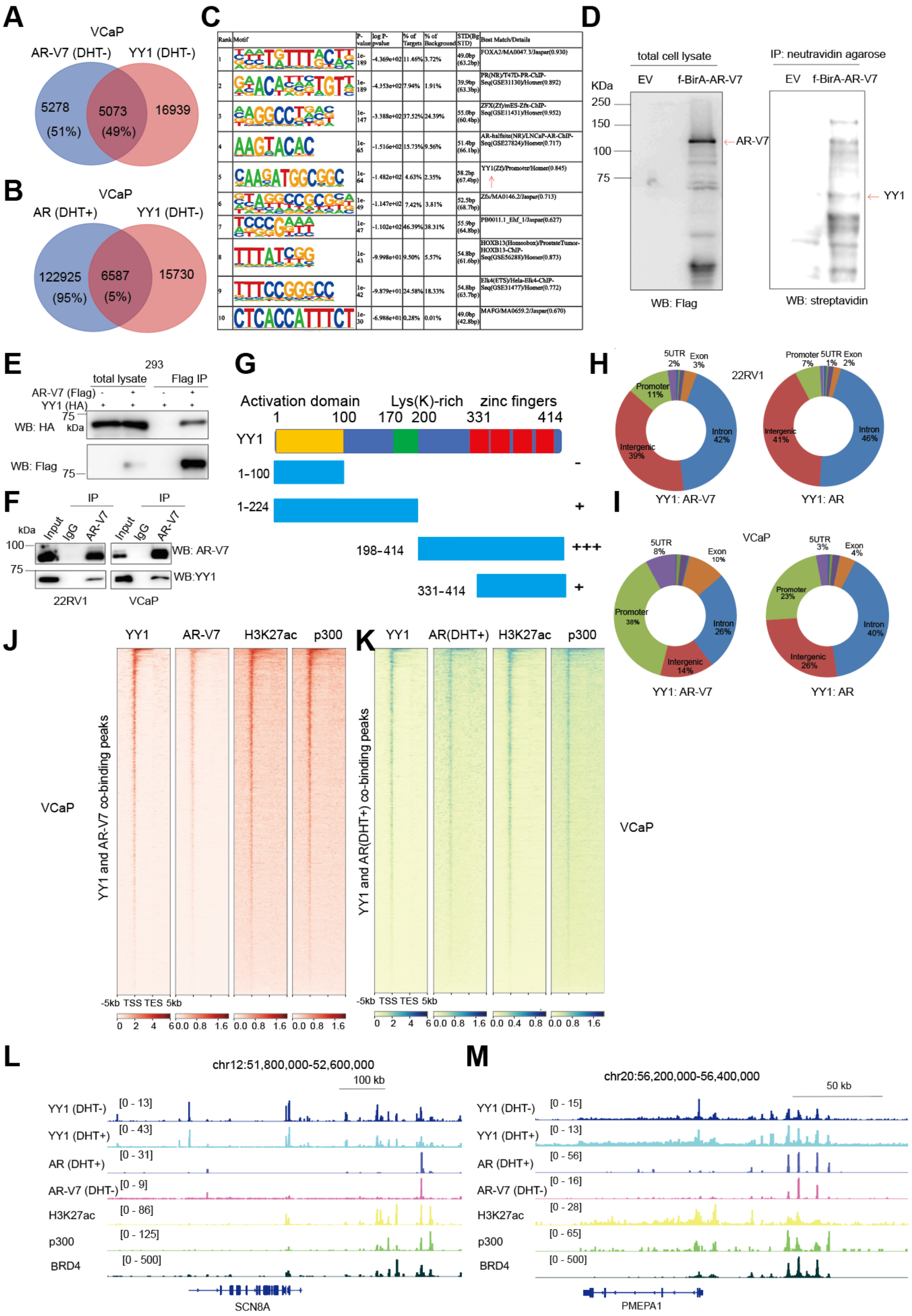


| Rank | Motif | P-value | log P-pvalue | % of Targets | % of Background | STD(Bg STD) | Best Match/Details |
|---|---|---|---|---|---|---|---|
| 1 | [illegible] | 1e-189 | -4.369e+02 | 11.46% | 3.72% | 49.0bp (63.2bp) | FOXA2/MA0047.3/Jaspar(0.930) |
| 2 | [illegible] | 1e-189 | -4.353e+02 | 7.94% | 1.91% | 39.9bp (63.3bp) | PR(NR)/T47D-PR-ChIP-Seq(GSE31130)/Homer(0.892) |
| 3 | [illegible] | 1e-147 | -3.388e+02 | 37.52% | 24.39% | 55.0bp (60.4bp) | ZFX(Zf)/mES-Zfx-ChIP-Seq(GSE11431)/Homer(0.952) |
| 4 | AAGTACAC | 1e-65 | -1.516e+02 | 15.73% | 9.56% | 51.4bp (66.1bp) | AR-halfsite(NR)/LNCaP-AR-ChIP-Seq(GSE27824)/Homer(0.717) |
| 5 | CAAGATGGCGGC | 1e-64 | -1.482e+02 | 4.63% | 2.35% | 58.2bp (67.4bp) | YY1(Zf)/Promoter/Homer(0.845) |
| 6 | [illegible] | 1e-49 | -1.147e+02 | 7.42% | 3.81% | 52.5bp (68.7bp) | Zfx/MA0146.2/Jaspar(0.713) |
| 7 | TCCCGAAA | 1e-47 | -1.102e+02 | 46.39% | 38.31% | 55.9bp (64.8bp) | PB0011.1_Ehf_1/Jaspar(0.627) |
| 8 | TTTATCGG | 1e-43 | -9.998e+01 | 9.50% | 5.57% | 54.8bp (61.6bp) | HOXB13(Homeobox)/ProstateTumor-HOXB13-ChIP-Seq(GSE56288)/Homer(0.873) |
| 9 | TTTCCGGGCC | 1e-42 | -9.879e+01 | 24.58% | 18.33% | 54.8bp (63.7bp) | Elk4(ETS)/Hela-Elk4-ChIP-Seq(GSE31477)/Homer(0.772) |
| 10 | CTCACCATTTCT | 1e-30 | -6.988e+01 | 0.28% | 0.01% | 49.0bp (42.8bp) | MAFG/MA0659.2/Jaspar(0.670) |

**Supplementary Fig. 6. YY1:p300 interaction acts to promote the AR/AR-V7-related oncogenic signaling in CRPC, conferring therapeutic resistance.**

**(A-B)** Venn diagram showing the overlap of YY1:AR-V7-cobound (**A**) or YY1:AR-cobound (**B**) peaks in VCaP cells. AR binding was stimulated with its ligand, DHT, prior to the cell collection.
**(C)** Homer-based unbiased motif search identified the YY1 consensus motif among the AR-V7 binding peaks in 22Rv1 cells.
**(D)** WB of the biotinylated proteins using a streptavidin-HRP conjugate antibody in 22Rv1 cells, which were stably transduced with a flag-tagged BirA* only (mock control [EV], lane 1) or BirA*-AR-V7 fusion (lane 2) and then treated with 50 uM of biotin for 24 h.
**(E)** Co-IP for AR-V7 interaction with YY1 in HEK293T cells, transfected with the indicated expression plasmids and then subjected to anti-Flag IP.
**(F)** Co-IP for interaction between the endogenous AR-V7 and YY1 in 22Rv1 (left) and VCaP (right) cells. Cells were subjected to anti-AR-V7 IP.
**(G)** Scheme showing a set of YY1 serial deletion constructs used for assessing YY1 interaction with AR-V7. CoIP results were summarized on the right, with "-", "+" and "+++" indicating no obvious interaction, weak and strong interaction, respectively.
**(H-I)** Pie chart showing genomic annotation of YY1:AR-V7-cobound (left) or YY1:AR-cobound (right) CUT&Tag peaks in 22Rv1 (**H**) and VCaP (**I**) cells.
**(J-K)** Heatmap showing the genomic binding signals of the indicated protein, ± 5 kb from the centers of either YY1:AR-V7-cobound (**J**) or YY1:AR-cobound (**K**) peaks identified in VCaP cells.
(**L-M**) IGV image of enrichment for the indicated antibodies at SCN8A (L) and PMEPA1 (M) in 22Rv1 cells.

Fig S7 Xu et al

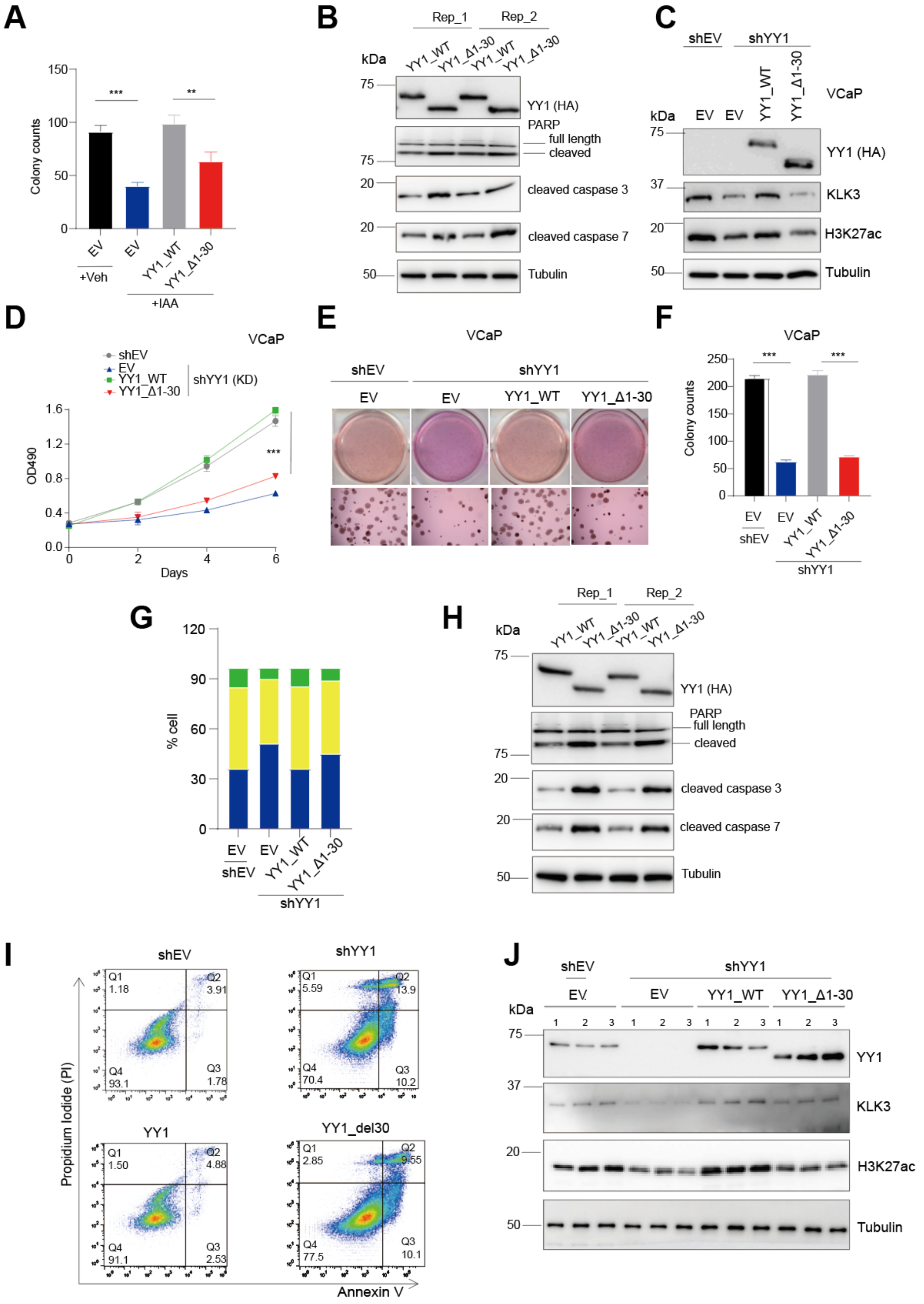


**Supplementary Fig. 7. The YY1$^{1\text{-}30aa}$-mediated p300 activation is critically involved in prostate tumorigenesis *in vitro* and *in vivo*.**

**(A)** Quantification of number of colonies formed by the 22Rv1 cells carrying the YY1-mAID KI alleles, treated with vehicle (Veh) or 5-Ph-IAA (IAA for YY1 degradation) and then rescued with either vector mock (EV), WT YY1, and YY1_Δ1-30aa (n = 3 replicates per group). *$P < 0.05$; **$P < 0.01$; ***$P < 0.001$.
**(B)** WB of the indicated apoptotic marker in the YY1-degraded 22Rv1 cells, rescued with WT YY1 and YY1_Δ1-30aa.
**(C-I)** WB of the indicated protein (**C**), kinetics of cell proliferation (**D**), the soft agar-based colony formation (**E**) and colony number quantification (**F**), analysis of cell cycle progression **(G)**, WB of various apoptotic markers **(H),** as well as representative profiles of Annexin-V and Propidium Iodide (PI)staining **(I)** using VCaP cells, which were stably transduced with control mock (shEV) or YY1-targeting shRNA (shYY1 for mediating YY1 KD), as well as the latter shYY1-expressing cells pre-rescued with either empty vector control (EV), WT YY1, or the YY1_Δ1-30aa (n = 3 replicates per group). **$P < 0.01$; ***$P < 0.001$.
**(J)** WB for the indicated protein using the 22Rv1 tumor xenografts collected at the end of study shown in **Fig. 7J-K**.
