## Supplementary material for "YY1 relieves p300 autoinhibition to promote histone acetylation in advanced prostate cancer, enhancing the oncogenic signaling of Androgen Receptor Splice Variant 7": Methods

### STAR METHODS

Detailed methods are provided in the online version of this paper and include the following:

- KEY RESOURCES TABLE
- RESOURCE AVAILABILITY
  - Lead contact
  - Materials availability
- EXPERIMENTAL MODEL AND SUBJECT DETAILS
  - Cell lines
  - Bacterial strains
- METHODS DETAILS
  - Antibodies and reagents
  - Plasmids
  - Co-Immunoprecipitation (Co-IP)
  - Stable and transient RNA interference
  - Virus Production and stable cell line generation
  - Colony formation assays
  - Isothermal titration calorimetry (ITC)
  - HAT assay
  - Luciferase reporter assays
  - Real-time Reverse transcription followed by quantitative polymerase chain reaction (RT-qPCR)
  - BioID
  - Gene Set Enrichment Analysis (GSEA)
  - CRISPR-Cas9-based editing of YY1
  - RNA-seq and data analysis
  - CUT&Tag and data analysis
  - In vivo tumor growth in xenograft models
- QUANTIFICATION AND STATISTICAL ANALYSIS

### STAR METHODS

### KEY RESOURCES TABLE

| REAGENT or RESOURCE | SOURCE | IDENTIFIER |
|---|---|---|
| Antibodies | | |
| FLAG® M2 antibody | Millipore Sigma | Cat # F1804; RRID:AB_262044 |
| YY1 (mouse polyclonal) | Santa Cruz | Cat # SC-7341; RRID: AB_2257497 |
| AR (N20) | Santa Cruz | Cat # SC-816; RRID: AB_1563391 |
| P300 | Cell Signaling Technology | Cat # 54062S; RRID: AB_2799450 |
| P300-Ac | Cell Signaling Technology | Cat # 4771S; RRID: AB_2262406 |
| FOXA1 | Abcam | Cat # ab23738; RRID: AB_2104842 |

| | | |
|---|---|---|
| Tip60 | Abcam | Cat # ab23886;<br>RRID: AB_778485 |
| BRD4 | Bethyl | Cat # A301-985A100;<br>RRID: AB_2620184 |
| PKFP | Cell Signaling Technology | Cat # 5412S;<br>RRID: AB_10692671 |
| Streptavidin | Cell Signaling Technology | Cat # 3999S;<br>RRID: AB_10544694 |
| H3K9ac | Cell Signaling Technology | Cat # ; 9649S<br>RRID: AB_10891758 |
| H3K14ac | Upstate | Cat #; 06-911<br>RRID: AB_310443 |
| H3K18ac | Active Motif | Cat #; 39129<br>RRID: AB_2561015 |
| H3K27ac | Abcam | Cat # ab4729;<br>RRID: AB_2118291 |
| H3K36ac | Active Motif | Cat #; 39379<br>RRID: AB_2561016 |
| H3K56ac | Upstate | Cat #; 07-677<br>RRID: AB_310307 |
| H4K16ac | Active Motif | Cat #; 61529<br>RRID: AB_2614970 |
| pan-aceylated H4 | Active Motif | Cat #;39925<br>RRID: AB_2561017 |
| H3K27me3 | Cell Signaling Technology | Cat # 9733S;<br>RRID: AB_2616029 |
| Cleaved caspase-3 | Cell Signaling Technology | Cat # 9661;<br>RRID:AB_2341188 |
| Cleaved caspase-7 | Cell Signaling Technology | Cat # 8438;<br>RRID:AB_11178377 |
| α-Tubulin | Cell Signaling Technology | Cat # 2144;<br>RRID:AB_2210548 |
| GAPDH | Cell Signaling Technology | Cat # 5174;<br>RRID:AB_10622025 |
| pan H3 | Cell Signaling Technology | Cat #; 4499S<br>RRID: AB_10544537 |
| Vinculin | Cell Signaling Technology | Cat # 9498;<br>RRID:AB_2797707 |
| Normal rabbit IgG | Cell Signaling Technology | Cat # 2729;<br>RRID:AB_1031062 |
| Anti-mouse IgG, HRP-linked antibody | Cell Signaling Technology | Cat # 7076;<br>RRID:AB_330924 |
| Anti-rabbit IgG, HRP-linked antibody | Cell Signaling Technology | Cat # 7074;<br>RRID:AB_2099233 |
| Bacterial and Virus Strains | | |
| DH5a competent cells | Thermo Fisher Scientific | Cat# 18265017 |
| One Shot Stbl3 competent cells | Thermo Fisher Scientific | Cat# C737303 |
| Chemicals, Peptides, and Recombinant Proteins | | |

| | | |
|---|---|---|
| 5-Ph-IAA | MedChemExpress | Cat # HY-134653 |
| Enzalutamide | Selleckchem | Cat # S1250 |
| Recombinant YY1 protein | Active Motif | Cat # 81119 |
| Recombinant p300 protein | Active Motif | Cat # 81158 |
| Critical Commercial Assays | | |
| RNeasy Plus Mini Kit (250) | Qiagen | Cat # 74136 |
| Dual-Luciferase® Reporter Assay System | Promega | Cat # E1960 |
| iSCRIPT cDNA Synthesis KIT | Biorad | Cat # 1708891 |
| iTaq Universal SYBR Green Supermix | Biorad | Cat # 1725125 |
| MycoAlert™ PLUS Mycoplasma Detection Kit | Lonza | Cat # LT27-286 |
| Lipofectamine 3000 Transfection Reagent | Thermo Fisher | Cat # L3000150 |
| NEBNext Ultra II RNA library Prep kit | New England Biolabs | Cat # E7770 |
| NextSeq™ 1000/2000 P2 XLEAP-SBS™ Reagent Kit | Illumina | Cat # 20100987 |
| NEBNext Multiplex Oligos for Illumina | New England Biolabs | Cat # E7335S |
| Annexin V-FITC Apoptosis Detection Kit | BD Biosciences | Cat # 556570; RRID:AB_2869085 |
| Monoclonal Anti-HA Agarose beads | Millipore Sigma | Cat # A2095; RRID: AB_257974 |
| SureBeads™ immunoprecipitation Kit with protein A and G conjugated magnetic beads | Biorad | Cat # 161-4833 |
| Deposited Data | | |
| Raw and analyzed mass spectrometry proteomics data | This study | PXD021901 |
| Raw and analyzed dataset of RNA-seq | This study | GEO: GSE158296 |
| Raw and analyzed dataset of CUT&Tag | This study | GEO: GSE158296 |
| Experimental Models: Cell Lines | | |
| 22Rv1 | ATCC | Cat # CCL-2505; RRID: CVCL_1045 |
| VCaP | ATCC | Cat # CRL-2876; RRID: CVCL_2235 |
| 293FT | Thermo Fisher | Cat # R70007; RRID: CVCL_6911 |
| Oligonucleotides | | |
| RT-qPCR oligos and sgRNAs used | This study (information in a supplemental table) | NA |
| Recombinant DNA | | |
| pCDH-EF1 with Flag-tagged YY1_Δ1-100 amino acids (aa) | This study | NA |
| pCDH-EF1 with Flag-tagged YY1_Δ1-30aa | This study | NA |
| pCDH-EF1 with HA-tagged YY1_AD_mutants #1-6 | This study | NA |
| pCDH-EF1 with HA-tagged YY1_AD_mutant (3E-3R) | This study | NA |
| pCDH-EF1 with HA-tagged YY1_AD_mutant (IL-SS) | This study | NA |
| pCDH-EF1 with HA-tagged YY1_AD_5aa-mutant | This study | NA |
| pCDH-EF1 with HA-tagged YY1_1-224aa | This study | NA |
| pCDH-EF1 with HA-tagged YY1_198-414aa | This study | NA |
| pCDH-EF1 with HA-tagged YY1_331-414aa | This study | NA |

| | | |
|---|---|---|
| pCDNA with Flag-tagged AR-V7 | This study | NA |
| pCDH-EF1 with HA-tagged p300_WT | This study | NA |
| pCDH-EF1 with Flag-p300_ΔCH3 (Δ1665-1809aa) | This study | NA |
| pCMV-Flag-tagged p300_WT | Gift from Dr. Lemasson | NA |
| pCMV-Flag-tagged p300_ΔHAT(Δ1472-1522aa) | Gift from Dr. Lemasson | NA |
| Software and Algorithms | | |
| GraphPad Prism 10 | GraphPad Software, LLC | Version 10.2.1 |
| FlowJo | FlowJo LLc | https://www.flowjo.com/ |
| ImageJ | ImageJ | https://imagej.nih.gov/ij/index.html |
| GSEA | [1] | https://software.broadinstitute.org/gsea/index.jsp |
| Homer | Heinz et al | Version 4.10.0/ http://homer.ucsd.edu/homer/ |
| IGV Browser | Robinson et al | https://software.broadinstitute.org/software/igv/ |
| Other | | |

### RESOURCE AVAILABILITY

#### Lead contact

#### Materials availability

All of the reagents reported in this study are available from the lead or correspondence contact with Materials Transfer Agreement as long as stocks remain available.

### EXPERIMENTAL MODEL AND SUBJECT DETAILS

#### Cell lines

HEK293T cells (purchased from ATCC) were maintained in the DMEM base medium supplemented with 10% of fetal bovine serum (FBS) and 1% of penicillin and streptomycin. Human prostate cancer lines, 22Rv1 and VCaP cells (ATCC), were cultured in the RPMI-1640 and DMEM base medium, respectively, supplemented with 10% of FBS and 1% of antibiotics. For treatment assays, cells were cultured under the ligand-deprived conditions for 3 days in phenol red-free RPMI-1640 medium containing 10% of charcoal-stripped serum, followed by treatment with vehicle, dihydrotestosterone (DHT), or DHT in combination with experimental compounds. Cells were incubated at 37°C in a humidified atmosphere with 5% of $CO_2$. Cell line identity was verified by short tandem repeat (STR) profiling, with mycoplasma contamination routinely monitored by using the MycoAlert™ PLUS Detection Kit (Lonza, Cat# LT27-286).

#### Bacterial strains

DH5α and Stabl3 chemically competent bacterial cells (Thermo Fisher Scientific) were used for plasmid transformation and propagation according to the manufacturer's instructions.

### METHOD DETAILS

#### Animal usage

All animal experiments were conducted in accordance with the protocol approved by the Institutional Animal Care and Use Committee (IACUC) of Duke University (Protocol #A254-23-12).

#### Plasmids.

A pFETCh[2]-based genomic targeting vector carrying a mAID-3×Flag-P2A-Neomycin (Neo) cassette was acquired from Addgene (cat 63934), amplified, and then inserted with the two homology arms flanking the stop codon of YY1 as described[2]. Additionally, the CRISPR guide RNAs (gRNAs) were designed using an online tool (https://chopchop.cbu.uib.no/) to target genomic sites near the stop codon of YY1 (sgRNA sequences are listed in Supplementary Table 3), synthesized, and were cloned into the Cas9-containing pSpCas9_BB_2A-GFP (PX458) plasmid (Addgene, 48138). This PX458 plasmid expressing YY1 sgRNA, together with the above pFETCh-YY1-mAID-3×Flag targeting donor vector, was used for creating cells with the knock-in (KI) alleles of YY1-mAID-3×Flag.

The following mammalian expression constructs were generated using the pCDH-EF1α-MCS-IRES-Puro/Neo lentiviral vector (System Biosciences, CD532A-2 and CD533A-2), which contains the Flag-tagged YY1, either wildtype (WT) or with the N-terminal amino acids (aa) truncated (Δ1-100aa and Δ1-30aa); HA-tagged YY1 with its AD mutated, including six 1-100aa mutants described in the paper (mutants #1–6) and 3 substitution mutants at YY1's AD: namely, a three-acidic-residues mutant of YY1's AD (E19R/E22R/E25R; termed as 3E-3R), a two-hydrophobic-residues mutant (I20S/L23S; termed as IL-SS), and a 5aa-mutant (a YY1_AD compound mutant harboring 3E-3R and IL-SS both); as well as the serial deletion constructs of HA-tagged YY1 encoding its sequence of 1-224, 198-414 or 331-414 aa. Additional constructs included the pcDNA-based expression vector carrying Flag-tagged AR-V7, as well as pCDH-EF1 vectors with HA-tagged p300 (HA-p300_WT) or Flag-tagged p300, either WT or with its CH3 domain deleted (Flag-p300_WT and Flag-p300_Δ1665–1809aa). Plasmids encoding a CMV-driven expression of Flag-tagged p300, either WT or with its HAT domain deleted (Flag-p300_WT and Flag-p300_ΔHAT [Δ1472–1522aa]), were kindly provided by Dr. I Lemasson (East Carolina University). For p300 BioID assay, A BirA* cDNA sequence was inserted in-frame to the N terminus of full-length p300 cDNA in the above generated pCDH-EF1α-MCS-IRES-Puro lentiviral construct. All plasmids were verified by Sanger sequencing before usage.

**Virus production and stable cell line generation.**
Stable knockdown (KD) cell lines were generated using the pLKO.1 shRNA lentiviral system following the manufacturer's protocol, as previously described[3]. Briefly, shRNA plasmids were co-transfected with the packaging vectors, psPAX2 and VSV-G, into 293FT cells using the polyethylenimine (PEI) transfection reagent. Viral supernatants were collected 48 hours post-transfection, filtered through a 0.45 μm filter, and used to infect target cells in the presence of 8 μg/mL polybrene (Santa Cruz Biotechnology, Cat# sc-134220). Infected cells were selected with the appropriate antibiotics to establish stable lines.

**Stable and transient RNA interference.**
The sgRNA sequences targeting YY1 were designed, synthesized and then cloned into the pLenti LRG-2.1_Neo vector (Addgene, cat#125593). A doxycycline (dox)-inducible lentiviral vector with SpCas9 was obtained from Dr. D Sabatini. The pLKO.1-based lentiviral shRNA vectors targeting the human YY1 (TRCN0000019894) were purchased from Sigma-Aldrich. All plasmid constructs were verified by Sanger sequencing. ON-TARGETplus SMARTpool siRNAs targeting YY1 (Cat# L-011796-00-0010), AR (Cat# L-003400-00-0010), FOXA1 (Cat# L-010319-00-0010) and non-targeting control siRNA (Cat# D-001810-10-05) were purchased from Dharmacon RNAi Technologies. Prostate cancer cells with either stable or transient gene KD were generated and used as described before[3,4].

**CRISPR–Cas9-based editing of YY1 and mAID-based YY1 degradation.**
A previously described approach termed pFETCh[2] was utilized to introduce a mAID with 3×Flag tag in-frame to the C-terminus of YY1 in 22Rv1 cells. To enable auxin-inducible degradation, we first generated a stable 22Rv1 cell line expressing OsTIR1(F74G)-9Myc using the lentiviral plasmid pLX_lenti_osTIR1_9Myc_P2A_Bsd (Addgene, 129717), in which the OsTIR1 coding sequence was mutated to carry the F74G substitution as previously described[5]. Lentiviral transduction and blasticidin selection were performed to establish a stable OsTIR1(F74G)-expressing background for subsequent AID-tagging experiments. For mAID knock-in (KI), two homology arms (HOM1 and HOM2) located before and after the stop codon of YY1 were synthesized as dsDNA genomic blocks, followed by cloning into a mAID-3×Flag-P2A-Neomycin (Neo) cassette-containing pFETCh donor vector (Addgene, 63934) using Gibson Assembly (New England BioLabs), yielding the pFETCh-YY1-AID-3×Flag targeting vector. After co-transfection of both the pFETCh donor and a sgRNA-expressing PX458 plasmid (targeting a region nearby the YY1 stop codon), cells were selected with 300 μg/ml G418 for 2 weeks. Drug-resistant cells were then plated into 96-well plates to derive single clones for genotyping. Direct Sanger sequencing using genomic DNA templates, along with anti-Flag WB, was used to confirm correct homologous recombination in the YY1-mAID-3×Flag KI lines, as we described before[6].

**Antibodies.**
Mouse antibodies against FLAG tag (M2; Cat # F1804), YY1 (Cat # SC-7341) and AR (N20; Cat # SC-816) were purchased from Santa Cruz Biotechnology. Antibodies against p300 (Cat # 54062S), acetylated-p300 (Cat # 4771S), PKFP (Cat # 5412S), Streptavidin

(Cat # 3999S), cleaved caspase 3 (Cat # 9661), cleaved caspase 7 (Cat # 8438), Tubulin (Cat # 2144), GAPDH (Cat # 5174), H3K27me3 (Cat # 9733S), general histone H3 (Cat # 4499S), Vinculin (Cat # 9498), normal rabbit IgG (Cat # 2729), anti-mouse IgG HRP-linked (Cat # 7076), and anti-rabbit IgG HRP-linked (Cat # 7074) were obtained from Cell Signaling Technology Inc. Rabbit antibodies against H3K9ac (Cat # 9649S), H3K14ac (Cat # 06-911), H3K18ac (Cat # 39129), H3K27ac (Cat # ab4729), H3K36ac (Cat # 39379), H3K56ac (Cat # 07-677), H4K16ac (Cat # 61529), and pan-acetyl-H4 (Cat # 39925) were purchased from Active Motif and/or Upstate. Anti-BRD4 antibody (Cat # A301-985A100) was purchased from Bethyl Laboratories, while anti-FOXA1 (Cat # ab23738) and anti-Tip60 (Cat # ab23886) antibodies were obtained from Abcam.

**Chemicals**
5-Ph-IAA (Cat # HY-134653) was purchased from MedChemExpress and enzalutamide (Cat # S1250) was obtained from Selleckchem. Usage of enzalutamide and DHT was described before[3].

**Recombinant protein**
The recombinant proteins of YY1 (Cat # 81119) and p300 (Cat # 81158) were purchased from Active Motif. The purified histone H3 was kindly provided by Dr. J Song (University of California Riverside), with the recombinant expression and purification reported before[7]. Protein containing the p300-TAZ2 domain and YY1's N-terminal 1–30aa was recombinantly expressed in bacterial cells (BL21 strain) and purified as described[8].

**Proximity-dependent biotin identification (BioID)**
A BirA* cDNA sequence[9,10] was fused in-frame with the N-terminus of the target protein (such as p300 and TF) within an MSCV-based retroviral vector, followed by virus production and generation of stable cell lines. BioID was performed as we previously described[11,12]. Briefly, cells were treated with 50 μM biotin for 24 hours, harvested from five 150-mm culture dishes, and washed twice with cold PBS. Cell pellets were resuspended in 1 mL of RIPA lysis buffer (10% glycerol, 25 mM Tris-HCl pH 8.0, 150 mM NaCl, 2 mM EDTA, 0.1% SDS, 1% NP-40, 0.2% sodium deoxycholate, supplemented with freshly added protease inhibitors). To digest nucleic acids, 1 μL of benzonase was added to the lysate, followed by incubation on ice for 1 hour. Lysates were clarified by centrifugation at maximum speed for 30 minutes at 4°C, and the resulting supernatant was incubated with 100 μL of NeutrAvidin agarose beads overnight at 4°C. Beads were sequentially washed twice with RIPA buffer and TAP lysis buffer (10% glycerol, 350 mM NaCl, 2 mM EDTA, 0.1% NP-40, 50 mM HEPES pH 8.0), followed by three washes with ABC buffer (50 mM ammonium bicarbonate, pH 8.0). Samples were then subjected to mass spectrometry analysis as described before.

**Co-immunoprecipitation (Co-IP)**
Co-IP was performed as previously described[3]. Briefly, cell pellets were lysed in the RIPA buffer freshly supplemented with 1× complete protease inhibitor cocktail (Roche) and PMSF. 1 milligram of total protein from the whole-cell lysates was incubated with the indicated antibodies overnight at 4°C on a rotator. Subsequently, 20 μL of protein G agarose beads (Roche, Cat# 11243233001) were added and incubated for an additional

2 hours at 4°C with rotation. Immune complexes were washed three times with 1m L of RIPA buffer, resuspended in 40 µL of 2× SDS loading buffer, and boiled at 90°C for 5 minutes. Samples were resolved by SDS-PAGE and transferred to PVDF membranes. Western blotting (WB) was conducted using standard procedures, and protein signals were visualized using enhanced chemiluminescence (ECL; GE Healthcare) according to the manufacturer's instructions.

**Colony formation assays.**

Cells were seeded in triplicate at a density of 20,000 cells per well in 6-well plates and cultured for 3 weeks, with fresh medium replaced twice weekly. Colonies were fixed and stained with iodonitrotetrazolium chloride solution (Sigma-Aldrich, Cat# I10406) to visualize colony formation.

**Isothermal Titration Calorimetry (ITC).**

Direct binding between YY1's N-terminal 1–30aa and the p300-TAZ2 domain was assessed using a MicroCal iTC200 system (Malvern) at 25 °C in buffer containing 100 mM Tris-HCl (pH 7.5) and 150 mM NaCl. Following an initial 0.4 µL priming injection, 13 successive injections of 500 µM YY1(1–30aa) were titrated into 300 µL of 50 µM pP300-TAZ2 protein in the sample cell, with constant stirring at 1,000 rpm. Binding isotherms were analyzed using a one-site binding model with MicroCal Origin 7.0 software (Malvern). Reported values represent the mean ± standard deviation from three independent experiments.

**Isothermal Titration Calorimetry (ITC).**

Direct binding between YY1's N-terminal 1–30aa and the p300-TAZ2 domain was assessed using a MicroCal iTC200 system (Malvern) at 25 °C in buffer containing 100 mM Tris-HCl (pH 7.5) and 150 mM NaCl. Following an initial 0.4 µL priming injection, 13 successive injections of 500 µM YY1(1–30aa) were titrated into 300 µL of 50 µM pP300-TAZ2 protein in the sample cell, with constant stirring at 1,000 rpm. Binding isotherms were analyzed using a one-site binding model with MicroCal Origin 7.0 software (Malvern). Reported values represent the mean ± standard deviation from three independent experiments.

**Histone Extraction**

Total histones were extracted with an acidic extraction protocol as previously described[13] and then subject to mass spectrometry-based quantification[14,15]. Briefly, Cell pellets were resuspended in nuclear isolation buffer (NIB; 15 mM Tris-HCl, pH 7.5, 60 mM KCl, 15 mM NaCl, 5 mM $MgCl_2$, 1 mM $CaCl_2$, 250 mM sucrose, 1 mM DTT) supplemented with protease and phosphatase inhibitor cocktail and 0.2% NP-40. The suspension was incubated on ice for 10 minutes, followed by centrifugation at 1,000 × g to isolate nuclei. The resulting nuclear pellets were washed twice with NP-40–free NIB to ensure complete removal of detergent. Isolated nuclei pellets were gently vortexed, and 0.4 N $H_2SO_4$ was added dropwise at a 5:1 volume ratio (e.g., 5 mL acid per 1 mL pellet) while vortexing. Samples were incubated on ice for at least 1 hour or up to overnight with intermittent mixing (e.g., 2 hours for ~1 million cells, longer for lower cell numbers). Following incubation, samples were centrifuged at 3,400 × g at 4°C, and the supernatant was collected. The extraction was repeated once, and both supernatants were pooled. To precipitate proteins, 100% trichloroacetic acid (TCA) was added to the pooled supernatant at 1:4 volume (final concentration: 20% TCA). The mixture was incubated on ice for ≥1 hour or overnight for dilute extracts, followed by centrifugation at 3,400 × g at 4°C. The resulting pellet was washed once with cold acetone containing 0.1% HCl and twice with 100% acetone (using only glassware for acetone handling). After centrifugation at 4,000 × g and air-drying, the histone pellet was resuspended in ~50 µL water. Histone purity was assessed by loading 5–10% of the extract on a 15% SDS-PAGE gel followed by Coomassie Blue staining. Protein concentration was determined by BCA or equivalent assay and adjusted to ~1 µg/µL. Extracts were snap-frozen in liquid nitrogen and stored or shipped on dry ice.

**HAT assay**

In vitro HAT assays were conducted in the 25 ml reaction mixture, which contained 0.5 ug of recombinant full-length p300, 1.0 ug of recombinant full-length YY1 and 10 mM of acetyl-coenzyme A in HAT assay buffer (50mM Tris–HCl, pH 8.0, 10% glycerol, 0.1mM EDTA and 1mM dithiothreitol), was incubated at 30C for 30 min, and then subjected to SDS–PAGE for protein separation. Separate HAT assay included additional 10 mg of the purified histone H3 (kindly provided by Dr. J Song, with the recombinant histone proteins described before[7]). Acetylation of histone H3 and p300 were analyzed by WB using antibody specific to acetyl-H3 (H3K27ac) and acetyl-p300 (p300K1499ac).

**Luciferase reporter assays.**

Cells were seeded in 24-well plates and transfected with the indicated plasmids. The pRL-CMV Renilla plasmid was used as an internal control. Luciferase activity was measured 48 h after transfection using the Dual-luciferase Reporter Assay System (Promega). Data were normalized to Renilla luciferase. Luciferase data are reported as mean ± s.e.m. of three independent experiments performed in triplicate.

**Real-time RT-qPCR.**

Total RNA was isolated using the RNeasy Mini Kit (Qiagen). First strand cDNA was synthesized using the High-Capacity cDNA Reverse Transcription Kit (Applied Biosystems). Real-time PCR was performed in triplicate using the iTaq Universal SYBR Green Supermix (Bio-Rad) and the Quant Studio 6 Flex Real-Time PCR System (Applied Biosystems). All values were normalized to internal controls (such as β-actin levels).

**RNA-seq and data analysis**

RNA was prepared as described before[3,11,16], using 2 million of the 22Rv1 or VCaP cells. Total RNA was extracted using the RNeasy Plus Mini Kit (Qiagen, 74136) according to the manufacturer's instructions, including an on-column DNase digestion step to remove genomic DNA. Polyadenylated RNA was enriched using the NEBNext Poly(A) mRNA Magnetic Isolation Module (New England BioLabs, E7490), followed by library construction with the NEBNext Ultra II RNA Library Prep Kit for Illumina (New England BioLabs, E7770 or E77705) following the vendor's protocol. Multiplexed libraries were sequenced on an Illumina NextSeq 500 platform using the NextSeq 550 High Output Kit v2.5 (Illumina, 20024906). For transcript quantification, reads were pseudo-aligned to the reference transcriptome using Salmon (v1.4.0). Alternatively, alignment was performed using MapSplice[17], and expression levels were quantified with RSEM[18]. Read counts were upper-quantile normalized, $log_2$-transformed, and used for differential expression analysis with DESeq2 (v1.38.2)[19].

### Gene ontology (GO) and Gene Set Enrichment Analysis (GSEA)

GO analysis was performed using Metascape[20] while GSEA was performed with the downloaded GSEA software (www.broadinstitute.org/gsea) by exploring the Molecular Signatures Database (www.broadinstitute.org/gsea/msigdb/annotate.jsp), as previously described[3,6,21,22].

### CUT&Tag and data analysis

CUT&Tag[23,24] was performed with modifications as we previously described[12,25]. Briefly, approximately 100,000 of human cancer cells (such as 22Rv1 and VCaP) were harvested per sample. For spike-in normalization, 5% of spike-in control cells (murine cells; see below) were added. Cells were washed once in Wash Buffer (20 mM HEPES pH 7.5, 150 mM NaCl, 0.5 mM Spermidine, 1× Protease Inhibitor Cocktail). Concanavalin A–coated magnetic beads (Bangs Laboratories, BP531) were activated in Beads Activation Buffer (20 mM HEPES pH 7.9, 10 mM KCl, 1 mM $CaCl_2$, 1 mM $MnCl_2$) and added to each sample (10 µL/sample), incubated at room temperature (RT) for 10 min. Bead-bound cells were resuspended in Digitonin 150 Buffer (20 mM HEPES pH 7.5, 150 mM NaCl, 0.5 mM Spermidine, 0.01% Digitonin, 1× Protease Inhibitor Cocktail) supplemented with 2 mM EDTA and the primary antibody (typically 1:50 dilution). For CUT&Tag of HA-tagged protein (such as rescued YY1) in 22Rv1 or VCaP cells, 5% of NIH-3T3 cells stably expressing H2B-GFP were added across samples as a spike-in normalization control and these assays used 1 µL of the primary antibody (HA or p300) plus 1 µL of anti-GFP antibody. For histone marks (H3K27ac, H3K27me3, H3K9ac) and YY1, the antibodies of which can recognize both human and mouse proteins, 5% of WT NIH-3T3 cells was included across all samples as an internal spike-in control for signal normalization and 1 µL of antibody against each one of the above factors was used.

Samples were rotated overnight at 4°C. The next day, samples were incubated with a secondary antibody (1:50 dilution in Digitonin 150 Buffer) for 30 min at RT. Unbound antibodies were removed by two washes with Digitonin 150 Buffer. The samples were incubated with a 1:200 dilution of pA-Tn5 adapter complex (pre-loaded with sequencing adapters) in Digitonin 300 Buffer (20 mM HEPES pH 7.5, 300 mM NaCl, 0.5 mM Spermidine, 0.01% Digitonin, 1× Protease Inhibitor Cocktail) for 1 hour at RT. Unbound enzyme was removed with two washes in Digitonin 300 Buffer. Tagmentation was initiated by resuspending the beads in Tagmentation Buffer (Digitonin 300 + 10 mM $MgCl_2$), followed by incubation at 37°C for 1 hour. Following removal of the tagmentation buffer, beads were incubated in 5 µL SDS Release Buffer (10 mM TAPS, pH 8.5; 0.1% SDS) at 58°C for 1 hour. Subsequently, 15 µL of SDS Quench Buffer (0.67% Triton X-100 in water) was added to neutralize SDS prior to PCR amplification. DNA was purified using Ampure XP beads per manufacturer's instructions and eluted in 10 mM Tris-HCl (pH 8.0). Indexed libraries were generated using a universal i5 primer and a barcoded i7 primer (unique per sample), using 2× PCR master mix. Libraries were cleaned with 0.9× Ampure XP beads and eluted in 30 µL Tris-HCl (10 mM, pH 8.0). Sequencing libraries were subjected to high-throughput paired-end sequencing using the Illumina NextSeq 550 or equivalent platform.

The resulting FASTQ files were aligned to both the main reference genome (GRCh37/hg19) and the spike-in control genome (GRCm38/mm10) using Bowtie2 (v2.3.5)[26]. Reads with non-primary alignments were removed using Samtools (v1.9), and PCR duplicates were marked and removed using Picard MarkDuplicates (v2.20.4). Blacklisted genomic regions were excluded using bedtools (v2.28.0). Peak calling was performed using MACS2 (macs2 callpeak -f BAMPE -g hs --keep-dup 1, v2.2.6)[27], and peak annotation was conducted using the annotatePeaks.pl function from HOMER[28]. Peak distribution relative to genomic features was also assessed using HOMER tools. For data visualization and signal profiling, deepTools (v3.3.0)[29] was used to generate normalized bigwig files (--normalizeUsing RPKM), heatmaps, and average signal profiles. Genome-wide comparisons between groups were visualized using the bamCompare function in deepTools. Read coverage and signal intensity at specific loci were visualized with the Integrative Genomics Viewer (IGV).

***In vivo* tumor growth in xenografted models.**

NSG mice were obtained from The Jackson Laboratory, housed in a specific pathogen–free (SPF) facility, and used as before[3,30]. In brief, a total of $1\times10^6$ 22Rv1 cells were resuspended in 100 µL of PBS containing 50% Phenol Red–free Matrigel (Corning, catalog no. 356237) and bilaterally subcutaneously (s.c.) injected into the dorsal flanks of castrated NOD/SCID/IL2Rγ-null (NSG) mice. Once the average tumor volume reached approximately 100 $mm^3$, mice were randomly assigned to vehicle, CBPD-268, or enzalutamide treatment cohorts. For in vivo administration, CBPD-268 was dissolved in 100% PEG200 and administered orally at a dose of 1 mg/kg/day, five days per week (Monday to Friday). Enzalutamide (MDV3100) was dissolved in DMSO and diluted in 1% carboxymethylcellulose (CMC; Sigma, cat# C4888) + 0.1% Tween 80 in water (final DMSO concentration 5%) to form a white slurry, which was homogenized by mixing or brief sonication prior to administration. The drug was delivered via oral gavage at a dose of 10 mg/kg/day. Tumor growth was monitored twice weekly using calipers, and tumor volume was calculated using the formula: $\text{length} \times \text{width}^2 \times 0.52$. Mice were sacrificed when tumors reached the maximum allowed size.

### <u>QUANTIFICATION AND STATISTICAL ANALYSIS</u>

Statistical analyses were performed using the GraphPad Prism version 9 software. To determine statistical significance, unpaired two-tailed Student's t test, one-way ANOVA, or the log-rank test was performed. Data are presented as mean ± SDs/SEMs from at least three independent experiments. *, **, and *** denote the *P* value of <0.05, 0.01 and 0.001, respectively. NS denotes not significant. As for CUT&Tag and RNA-seq datasets, the methods used for statistical analysis are described in the above sections.

### SUPPLEMENTAL INFORMATION

Supplemental information can be found online at
PDF containing Figure S1-S7

### ACKNOWLEDGEMENTS

We thank all members of the Cai and Wang laboratories for their helpful discussions. This work was supported in part by the U.S. National Institutes of Health (NIH) grant R01CA262903 (to L.C.) and institutional funds from Duke University (to G.G.W. and L.C.). Core facilities affiliated with the Duke Cancer Institute (DCI) are supported in part by the NCI Cancer Center Support Grant (P30CA014236) and the NIH Shared Instrumentation Grant (1S10RR027867-01) for confocal microscopy systems. Figures were created with BioRender.com.

### AUTHOR CONTRIBUTIONS

C.X. carried out a majority of the experiments. D.W. and K.J. performed AlphaFold analysis under the supervision of G.G.W. X.R. synthesized the p300/CBP degrader, CBPD-268, under the supervision of J.J. BioID proteomic analyses using mass spectrometry were conducted by S.G.M. and R.D.E., under the supervision of A.J.T, while the histone mass spectrometry studies were conducted by N.V.B, under the supervision of B.A.G.. RNA-seq and CUT&Tag data analyses were performed by C.X. and B.P., under the supervision of G.G.W. and L.C. X.L. conducted ITC assays. Data interpretation was carried out by C.X., D.W., G.G.W., and L.C. The project conceptualization and experimental designs were jointly supervised by G.G.W. and L.C.. The manuscript was written by C.X., G.G.W., and L.C., with input from all co-authors.

### DECLARATION OF INTERESTS

The authors declare no competing interests.
